# Functional Genomics of Gastrointestinal *Escherichia coli* Isolated from Patients with Cancer and Diarrhea

**DOI:** 10.1101/2023.05.31.543115

**Authors:** Hannah Carter, Justin Clark, Lily G. Carlin, Ellen Vaughan, Anubama Rajan, Adilene Olvera, Xiaomin Yu, Xi-Lei Zeng, Amal Kambal, Michael Holder, Xiang Qin, Richard A. Gibbs, Joseph F. Petrosino, Donna M. Muzny, Harsha Doddapaneni, Vipin K. Menon, Kristi L. Hoffman, Qingchang Meng, Matthew C. Ross, Sara J. Javornik Cregeen, Ginger Metcalf, Robert Jenq, Sarah Blutt, Mary K. Estes, TMC-GCID team, Anthony Maresso, Pablo C. Okhuysen

**Affiliations:** Department of Molecular Virology and Microbiology, Baylor College of Medicine, Houston, TX, USA; Department of Infectious Diseases, Infection Control and Employee Health, The University of Texas MD Anderson Cancer Center Houston, TX, USA; The Human Genome Sequencing Center, Baylor College of Medicine, Houston, TX, USA; Alkek Center for Metagenomics and Microbiome Research, Baylor College of Medicine, Houston, TX, USA; Department of Genomic Medicine – Clinical, The University of Texas MD Anderson Cancer Center Houston, TX, USA; Department of Medicine, Baylor College of Medicine, Houston, TX, USA; Section of Infectious Diseases, Baylor College of Medicine, Houston, TX, USA

**Keywords:** Diarrhea, Enteroaggregative, *E. coli*, Intestinal organoids, Colonoids, Cancer

## Abstract

We describe the epidemiology and clinical characteristics of 29 patients with cancer and diarrhea in whom Enteroaggregative *Escherichia coli* (EAEC) was initially identified by GI BioFire panel multiplex. *E. coli* strains were successfully isolated from fecal cultures in 14 of 29 patients. Six of the 14 strains were identified as EAEC and 8 belonged to other diverse *E. coli* groups of unknown pathogenesis. We investigated these strains by their adherence to human intestinal organoids, cytotoxic responses, antibiotic resistance profile, full sequencing of their genomes, and annotation of their functional virulome. Interestingly, we discovered novel and enhanced adherence and aggregative patterns for several diarrheagenic pathotypes that were not previously seen when co-cultured with immortalized cell lines. EAEC isolates displayed exceptional adherence and aggregation to human colonoids compared not only to diverse GI *E. coli*, but also compared to prototype strains of other diarrheagenic *E. coli*. Some of the diverse *E. coli* strains that could not be classified as a conventional pathotype also showed an enhanced aggregative and cytotoxic response. Notably, we found a high carriage rate of antibiotic resistance genes in both EAEC strains and diverse GI *E. coli* isolates and observed a positive correlation between adherence to colonoids and the number of metal acquisition genes carried in both EAEC and the diverse *E. coli* strains. This work indicates that *E. coli* from cancer patients constitute strains of remarkable pathotypic and genomic divergence, including strains of unknown disease etiology with unique virulomes. Future studies will allow for the opportunity to re-define *E. coli* pathotypes with greater diagnostic accuracy and into more clinically relevant groupings.

## INTRODUCTION

Infections are a serious threat and the leading cause of death in cancer patients.^1^ Among patients with hematological malignancies, bacterial infections are responsible for 43% of deaths and polymicrobial infections are responsible for an additional 11% of deaths^2^. Patients with cancer are at increased risk for infection due to neutropenia, immunosuppression, and other treatment-related complications^1^.

Furthermore, antibiotic resistance^3, 4^ limits treatment options and compromises their recovery. Chemotherapy, radiation, immunosuppression, hospitalization, and use of antibiotics^5^ all disrupt the microbial ecosystems of the intestinal tract and enables the increase in Enterobacteriaceae and *Enterococcus spp* abundance^6, 7^. Dominance by such pathobionts including distinct *E. coli* types place patients at risk for diarrhea and future infections due to translocation from the intestinal lumen.

Disease-causing *E. coli* have been categorized into “pathotypes” based on clinical syndromes, organs involved, adherence and aggregation patterns, and the presence of certain toxins. For some *E. coli* pathotypes -- for example, enterotoxigenic *E. coli* (ETEC) – the pathotype criteria describe a group of similar strains that share a common disease mechanism and clinical symptoms. However, enteroaggregative *E. coli* (EAEC), as currently defined, are a collection of heterogeneous strains lacking a unifying disease mechanism^8–12^.

In healthy individuals, infection with EAEC causes self-limited watery diarrhea with or without blood and mucus, abdominal pain, nausea, vomiting, and fever^13^. In children living in resource poor environments, EAEC can also cause subclinical asymptomatic infection resulting in chronic intestinal inflammation^14^ and is associated with impaired growth and development. Asymptomatic children and adults serve as carriers and potentially transmit to others. In volunteer challenge studies, not all EAEC strains can cause symptoms and not all individuals are susceptible^11^ adding the confounding factor of differential host-pathogen interactions^15^ that is likely associated with pathogen and host genetics^16–18^ and likely influenced by host immunity. Diarrheal disease is associated with infections of sequence type ST40 while asymptomatic carriage is higher among ST31 strains^8^, but the current pathotype system for describing EAEC does not allow stratification of strains based on their pathogenicity or host-specificity.

A multistep model has been proposed for EAEC pathogenesis beginning with bacterial auto-aggregation, adherence to the intestinal epithelium, and biofilm formation preceding the secretion of bacterial toxins and effectors responsible for host fluid secretion^9^. The EAEC pathotype was first described as fecal *E. coli* with the ability to form a unique “stacked brick” adherence pattern on HEp2 cells^19^ and EAEC were later identified to carry: the transcriptional regulator *aggR*, dispersin (*aatA*) and associated transporters and accessory genes, and a complete aggregative adherence fimbriae (AAF) cassette^20^. Studies using human intestinal enteroids (HIEs) suggest EAEC adherence to human intestinal cells is far more nuanced than previously believed: on HIE monolayers, EAEC forms localized 3-dimensional microcolonies and large aggregative networks in addition to the characteristic “stacked-brick” AA pattern^15^. However, EAEC’s “stacked brick” pattern^21^, aggregative adherence (AA) to Hep2 cells^22, 23^ and human colonoids^20^, and biofilms on abiotic surfaces^24, 25^ can be facilitated by other combinations of genes besides those currently used to identify EAEC. Such strains with these abilities but lacking *aggR*, *aata*, and an AAF cassette are named “atypical EAEC”^26^ and their pathogenicity is still contested. Remarkably, *aggR- Citrobacter freundii* strains able to form an AA pattern have also been isolated^21^ further complicating our understanding of the connection between this aggregative phenotype and clinical symptoms.

The EAEC pathotype as it is currently defined likely includes both pathogenic and non-pathogenic strains^27^ and aggregative adherence to Hep2 cells^19^ is likely an inaccurate predictor of adherence to human intestinal epithelium^8^ or human patients^20^. In patients with diarrhea where EAEC can be identified, it is generally assumed that EAEC is the etiological cause of disease. However, as EAEC can inhabit the GI tract without causing diarrhea,^28, 29^ it is possible that additional factors are contributing to the disease presentation in these patients. Here, we investigate *E. coli* strains isolated from cancer patients presenting with diarrhea and a stool sample positive for EAEC. In addition to EAEC strains, we isolated several enigmatic *E. coli* which we have characterized alongside typical EAEC isolates using intestinal organoids.

## RESULTS

### Isolation of *E. coli* from patients

Stool samples were collected from patients with cancer and diarrhea at The University of Texas MD Anderson Cancer Center and transported to the clinical microbiology laboratory in liquid Cary Blair transport media for diagnostic purposes and tested for the presence of enteric pathogens using the GI multiplex BioFire FilmArray® nucleic acid amplification test (NAAT) panel. EAEC was identified by NAAT in 29 patients (Figure 1). Residual stool was then plated onto MacConkey and EMB agar and 10 individual coliform colonies were isolated and stored in peptone agar stabs for further study. In 2018, coliforms were successfully isolated from storage stabs in 16/29 (55%) of patients and single colonies were selected by plating onto MacConkey and EMB agar plates for further characterization by MALDI TOF and whole genome sequencing (WGS). MALDI TOF demonstrated that the majority of isolates 15/16 (94%) were *E. coli* except for one which was reported as *Citrobacter freundii.* Upon examination of by WGS, genes characteristic of EAEC were identified in 6/15 (40%) isolates with the remaining 9 isolates (from 7 patients) belonging to diverse gastrointestinal *E. coli*. We then compared the clinical characteristics of patients in whom EAEC was identified by WGS to the group of patients in whom diverse *E. coli* were found, while also quantifying the ability of clinical *E. coli* isolates from both groups to adhere to human intestinal organoids when compared to adhesion phenotypes to reference prototypic diarrheagenic pathotypes.

**Figure 1.**
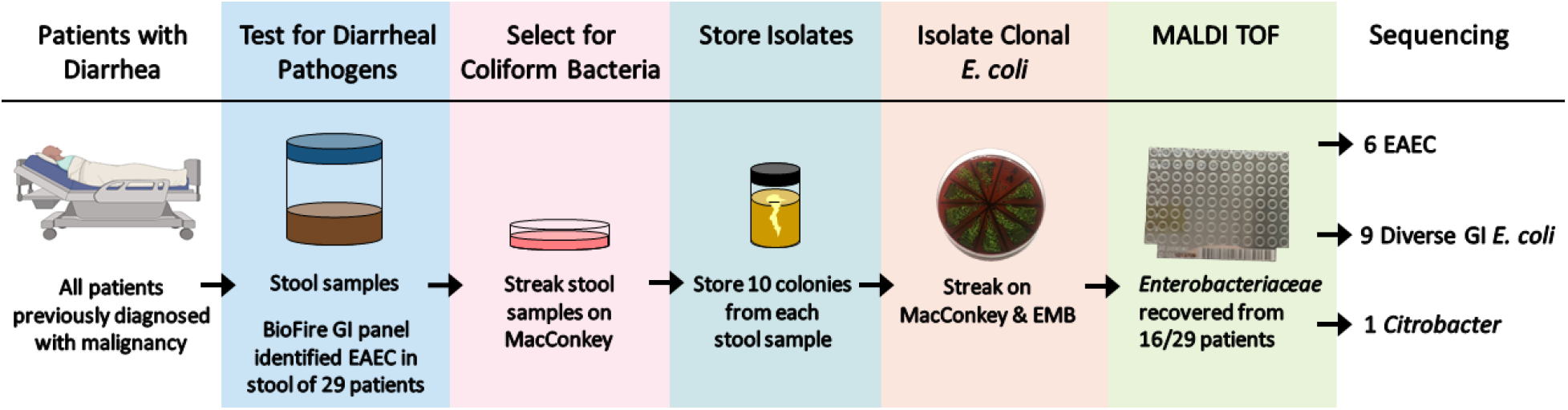
Isolation of Enterobacteriaceae strains in this study. Between 2015 and 2019, we collected stool samples from patients with diarrhea and EAEC (N=29) identified by the BioFire® FilmArray® Gastrointestinal (GI) in the clinical microbiology laboratory. Coliforms were grown in selective media and individual colonies stored as agar stabs in room temperature conditions to prevent plasmid loss. Individual strains were isolated with selective media for *Enterobacteriaceae* and passaged for clonal purity. The species of each isolate was confirmed with MALDI-TOF and whole genome sequencing.

Patient demographics and clinical characteristics were available in 28 of 29 cases and are shown in Table 1. NAAT identified EAEC in patients 17-81 years of age presenting with acute or chronic diarrhea. Both genders were identified in similar proportions. The majority of patients (18/28, 64%) had underlying hematologic malignancies including acute leukemia, lymphoma and multiple myeloma and 10 (36%) had solid tumors. Ten patients had undergone one or more hematopoietic stem cell transplantation (HSCT) and 4 presented with graft versus host disease (GvHD). Although the proportions of hematologic malignancies were similar in patients in whom *E. coli* could be recovered from those in whom it was not recovered, a subgroup analysis showed that the odds of recovering an *E. coli* isolate for characterization was significantly higher in patients who had undergone HSCT than those that had not (9/14, 64% vs. 1/14, 7%; p=0.004, OR 23. 4, 95% CI 2.32-235) and included all 4 patients with GVHD. Of note, several patients had the co-occurrence other bacterial enteric pathogens on BioFire NAAT but were not recovered in culture.

**Table 1.**
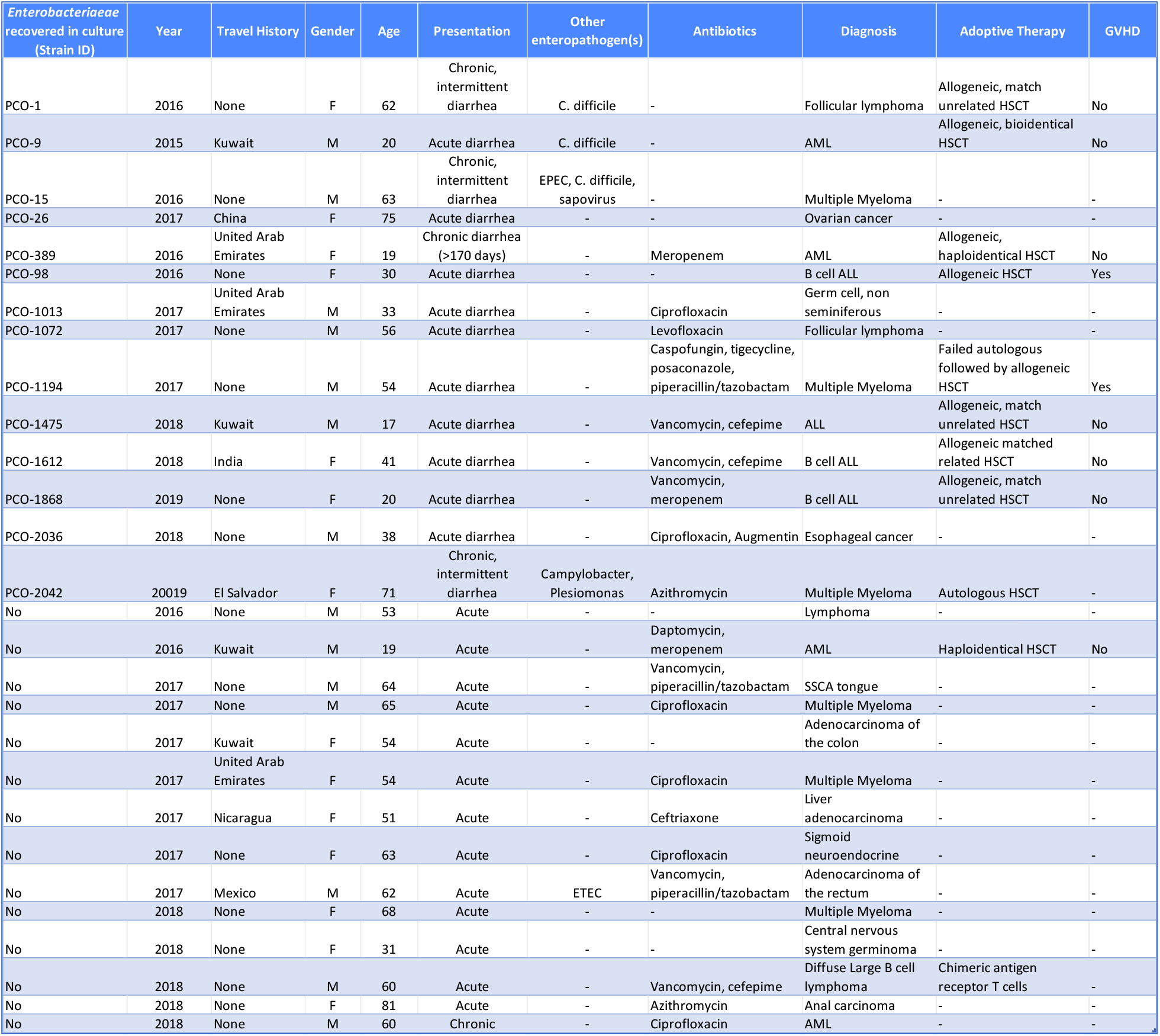
Clinical and epidemiology data from patients with cancer and diarrhea in whom EAEC was identified by NAAT and coliforms were recovered on subculture. Patients presented with acute and chronic diarrhea. Most (12/14 or 86%) were immunosuppressed and 11/14 or 79% were hematologic malignancies.

### Adherence of a Diverse Array of Gastrointestinal *E. coli* to Human Colonoids

To better understand the nature of the EAEC and the other diverse GI *E. coli* strains isolated from the above-referenced patients and how these compared to classic diarrheagenic *E. coli* pathotypes, we performed adherence and cytotoxic assays on transverse colonoids from patient 202 (TC202). For this, clinical isolates, prototype reference strains belonging to classical diarrheagenic *E. coli* pathotypes^30^ (Enteroinvasive (EIEC), Enterohemorrhagic (EHEC), Enteropathogenic (EPEC), and Enterotoxigenic (ETEC) *E. coli*), and a control strain (HS) which is devoid of all virulence genes characteristic of diarrheagenic *E. coli* and does not adhere to HEp-2 cells, were assessed for adherence and cytotoxicity in a transverse colon-derived HIE colonoid^31^ monolayer model of infection as previously established by our group^15, 32^. When adherence at 6 hours post infection was assessed, the control strain HS^33^, showed little adherence or aggregative ability (Figure 2 “Controls”). In contrast, the diarrheagenic strains demonstrated various adherence phenotypes. EIEC adhered predominantly as single bacteria, whereas ETEC, EHEC, and EPEC formed diverse and significant aggregates that mirrored recently and previously reported patterns characteristic of EAEC, including formation of microcolonies, sheet-like aggregative adherence, mesh-like aggregation, and network formation^15, 32^. To our knowledge, this is the first assessment of colon-derived HIEs that examines the adherence characteristics of EIEC, EHEC^34^, EPEC, and ETEC. We found that EPEC demonstrated its characteristic localized adherence pattern on colonoids and diffuse and network-like adherence as previously reported^32^. Unexpectedly, ETEC adhered in greater number than both EPEC and EIEC clustering together in a pattern previously unseen in adherence assays using immortalized cell lines.^35^

**Figure 2.**
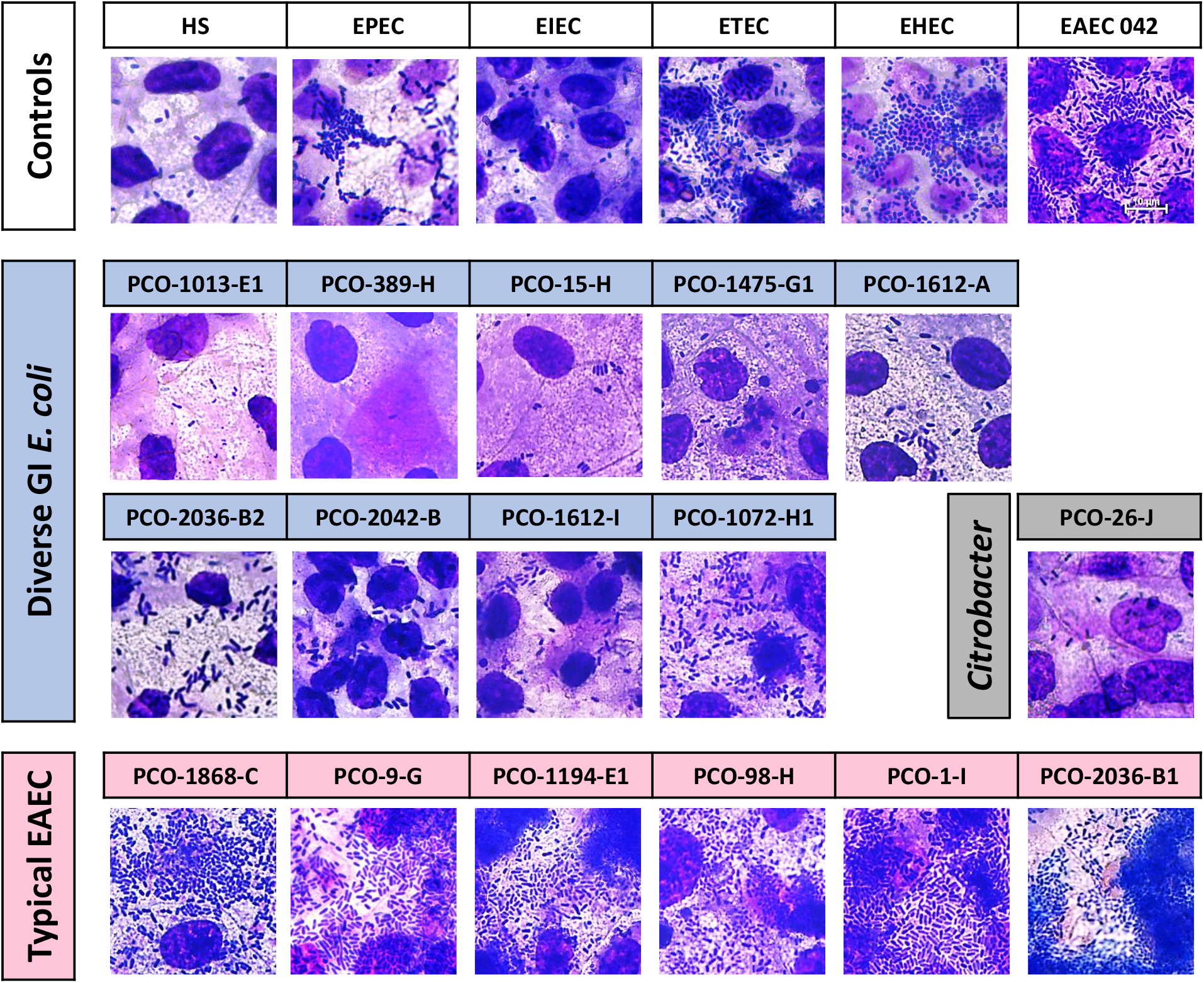
Adherence patterns of prototype diarrheagenic E. coli pathotypes and clinical Enterobacteriaceae isolates on human colonoids. HIE colonoid monolayers were infected with *E. coli* at an MOI = 10 and incubated for 6 hours. Media was removed and slides were fixed and stained using Hema 3 stain which is comparable to Wright-Giemsa stains. Slides were imaged at 100x magnification to gather representative images (scale bar shown in top right image). For each experiment, 5 images were taken per well across 4 separate wells of each condition. Each strain was tested in at least two independent experiments on separate days. Contrast and brightness adjusted for representative images above. Strains are presented in increasing order of the total number of adhered bacteria (as shown in Figure 3). Note that 1612-A and 1612-I came from the same patient, as well as 2036-B1 and 2036-B2. Neither of these pairs from the same patient sample are clonal.

Having established a reference pattern of adherence for known *E. coli* pathotypes in the colonoid HIE model, we turned our attention to carrying out a similar analysis with the clinical *E. coli* isolated from patients with cancer and diarrhea. We began with the strains deemed by WGS sequencing to carry *aggR* and therefore, categorized as typical EAEC^26, 36^. All six typical EAEC clinical isolates and the reference strain 042 demonstrated the characteristic stacked-brick adherence pattern in this assay, and predominately formed two-dimensional aggregative networks^15, 32^ spanning the majority of the colonoid monolayer (Figure 2 “typical EAEC”). While typical EAEC showed the characteristic aggregative adherence pattern, the diverse GI *E. coli* were quite varied in phenotype (Figure 2 “diverse GI *E. coli*”). *E. coli* isolates 1013-E1, 389-H, and 15-H adhered poorly to HIE colonoids and did so in a sparse, diffuse pattern. This phenotype also describes the *C. freundi* isolate, 26-J, which demonstrated mostly diffuse adherence. Isolates 1475-G1 and 1612-A mainly adhered in a diffuse pattern with the addition of small bacterial clusters. The most consistent phenotype observed was mesh-like aggregation interspersed with diffusely adhered singlet bacteria, as seen with isolates 2036-B2, 2042-B, 1612-I, and 1072-H1.

These four strains also demonstrated clustering of bacteria on the epithelial colonoid surface. Without additional genotyping of strain 1072-H (described later and depicted in Figure 5), 1072 would have been classified as an atypical EAEC based on adherence phenotype and absence of AggR.

To minimize ambiguity in our assessments, we applied a quantitative approach to measure the adherence and aggregation characteristics for each strain using quantitative light microscopy (Figure 3).

**Figure 3.**
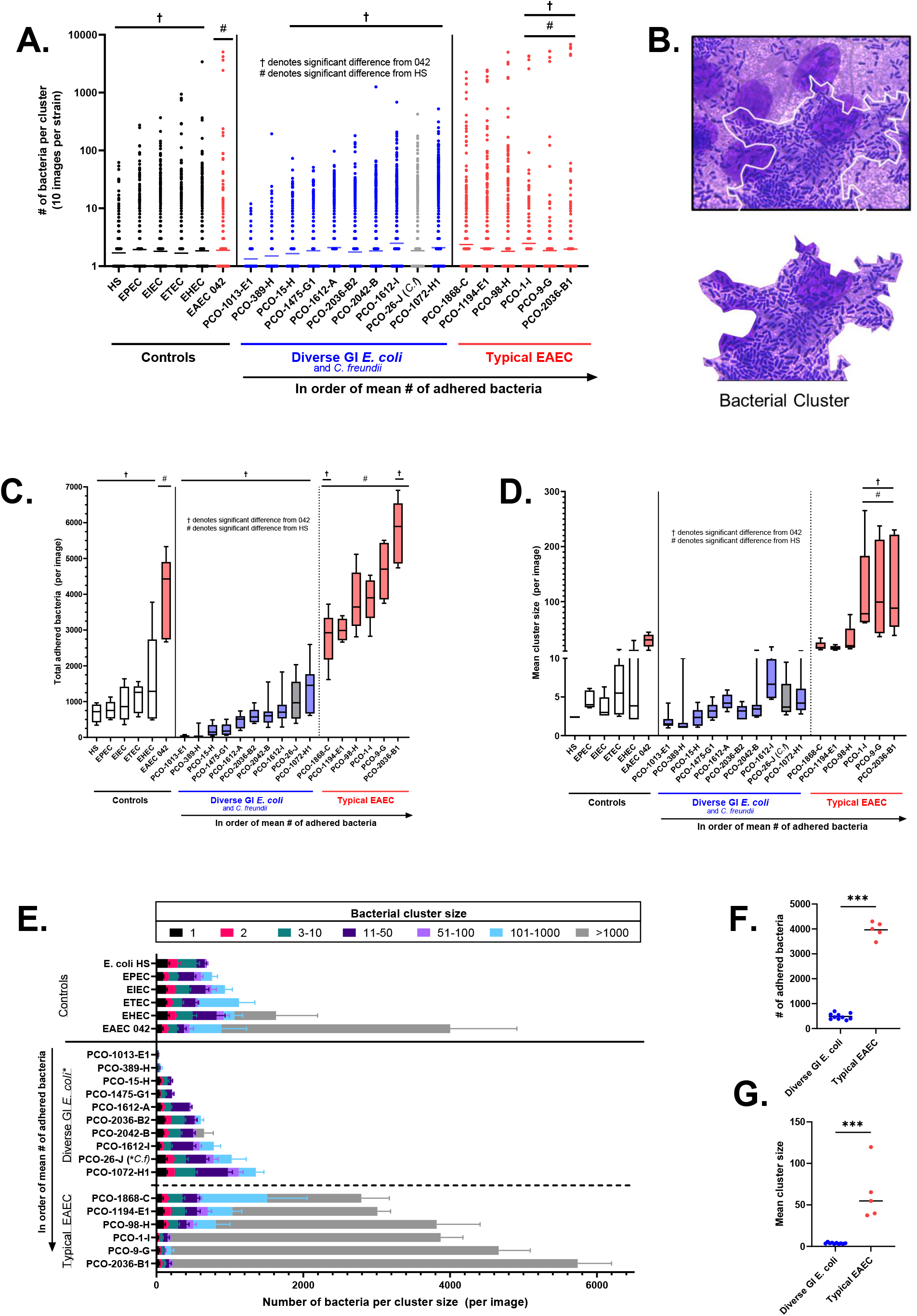
Adherence of prototype diarrheagenic E. coli and clinical Enterobacteriaceae isolates to human intestinal colonoids. From representative images, the total number of adhered bacteria and the number of bacteria in each bacterial cluster were counted following 6 hour infection and Hema3 staining (B). At least 5 images were counted manually for each strain. The number of bacteria in each cluster is depicted in (A) with every data point and the geometric mean shown. From this, the total number of adhered bacteria per image (C) and the mean cluster size (D) were calculated. These are shown with the minimum and maximum values as the ends of the box and whisker plots and the mean as the center line. Raw data was used to calculate the number of bacteria in each cluster size category allowing for the visualization depicted in (E). For each strain, the mean of each cluster size is plotted and error bars depict SEM. Comparisons were made using ordinary One-Way ANOVA with Dunnett’s multiple comparisons test. In (F) and (G) the typical EAEC strains were compared against the diverse GI *E. coli* using a Mann-Whitney U test.

We first determined the number of bacteria present in each bacterial cluster (Figures 3 A and B), then calculated the total number of adhered bacteria (Figure 3C) and finally, we determined the mean size of the clusters (Figure 3D). Throughout all figures, isolates are shown in increasing order of total adhered bacteria on the TC202 colonoids.

The number of bacteria in each and every cluster across five different images is shown in Figure 3A and is depicted as raw data allowing for visualization of adherence densities amongst strains. The non-pathogenic control strain HS exclusively formed small clusters with <100 bacteria while EPEC, EIEC, and ETEC all formed clusters holding between 100 and 10^3^ bacteria. Across the multiple images representing multiple wells, EHEC formed clusters with >10^3^ bacteria and typical EAEC strain 042 formed clusters ranging from 10^3^ to 10^4^ bacteria. The control strain HS and reference strains showed significantly fewer bacteria per cluster when compared to 042 (HS, EPEC, EIEC, ETEC, EHEC; p = 0.0072, p = 0.0321, p = 0.0051, p = 0.0147, p = 0.0138 respectively). Clinical EAEC isolates 98-H, 1194-E1, and 1868-C formed clusters with bacteria that were comparable to EAEC 042 and strains 1-I, 9-G, and 2036-B1 formed clusters with a larger number of bacteria per cluster than EAEC 042 (p < 0.0001, p = 0.0242, p < 0.0001 respectively).

We then examined the total number of adhered bacteria (Figure 3C) and the mean size of bacterial clusters formed (Figure 3D) on HIEs. In the case of EPEC, EIEC, and ETEC reference strains, the number of adhered bacteria was similar to control strain HS. In contrast, EHEC and EAEC 042, demonstrated a larger number of bacteria adhered to the HIE when compared to HS (p=0.0477 and p<0.001 respectively). EAEC 042 showed a significantly larger number of adherent bacteria than EPEC, EIEC, ETEC, EHEC, and non-pathogenic HS (p<0.0001, p<0.0001, p<0.0001, and p=0.0001) thereby setting EAEC apart as a pathotype with exceptional adherence ability (Figure 3C) to human colonoids. Among typical EAEC isolates, including the control strain 042, the aggregative networks formed frequently consisted of >1000 bacteria (Figure 3 C and E). Of note, several of the clinical typical EAEC strains from patients surpassed the aggregative adherence abilities of prototype strain 042. For example, strains 1-I, 9-G, and 2036-B1 displayed significantly increased mean cluster size (p<0.0001, p=0.0045, p<0.0001) and 2036-B1 showed significantly higher adherence (p<0.0001). This strain had the greatest mean cluster size observed in this study (Figure 3E) and the largest single cluster with 6826 bacteria aggregating together in a continuous group (Figure 3A).

Interestingly, the diverse GI *E. coli* isolates showed a range of aggregative phenotypes but with overall less total adherence than typical EAEC strains. Strains 1013-E1 and 389-H showed the poorest adherence to colonoids each with a mean of <100 adhered bacteria (per image) (Figure 3C). Isolates 15- H, 1475-G1, 1612-A, 2036-B2, and 2042-B had >100 adhered bacteria per image, but still had a total adherence less than that of control strain HS (Figure 3C). Of these diverse isolates, only 1612-I, 26-J (the *C. freundii* strain), and 1072-H1 showed a mean adherence greater than HS (Figure 3C). Of note, the only *Citrobacter* strain, 26-J, was second highest in adherence amongst the diverse GI isolates. Even the isolates best at adherence amongst non-EAEC strains, displayed markedly less adherence compared to typical EAEC strains. There is a sharp difference apparent between typical EAEC strains and our diverse GI *E. coli* group with a significant difference in both adhered bacteria (p=0.0007) and mean cluster size (p=0.0007) using a Mann-Whitney U test (Figure 3 F and G).

Amongst typical EAEC and diverse *E. coli*, total adherence and mean cluster size showed a strong positive correlation (Figure 4 B-D), but some isolates defied this trend. For example, in the diverse *E. coli* category, 1612-I had the greatest mean cluster size despite being bested in total adherence by 1072-H1. This discrepancy captures how strain 1612-I has an increased tendency to aggregate compared to strains with similar total adherence (such as 2036-B2). (These relationships are further explored in Figure 4 B-D). Surprisingly, strain 2042-B was uniquely able to form clusters of >10^3^ bacteria – a feature observed ubiquitously in EAEC strains in this study – despite 2042-B showing a similar number of adhered bacteria as other non-EAEC strains (Figure 3C). Most interestingly, each of the diverse GI *E. coli* strains had fewer total adhered bacteria compared to EAEC 042 (Figure 3C) but did not show significant differences in mean cluster size (Figure 3D).

**Figure 4.**
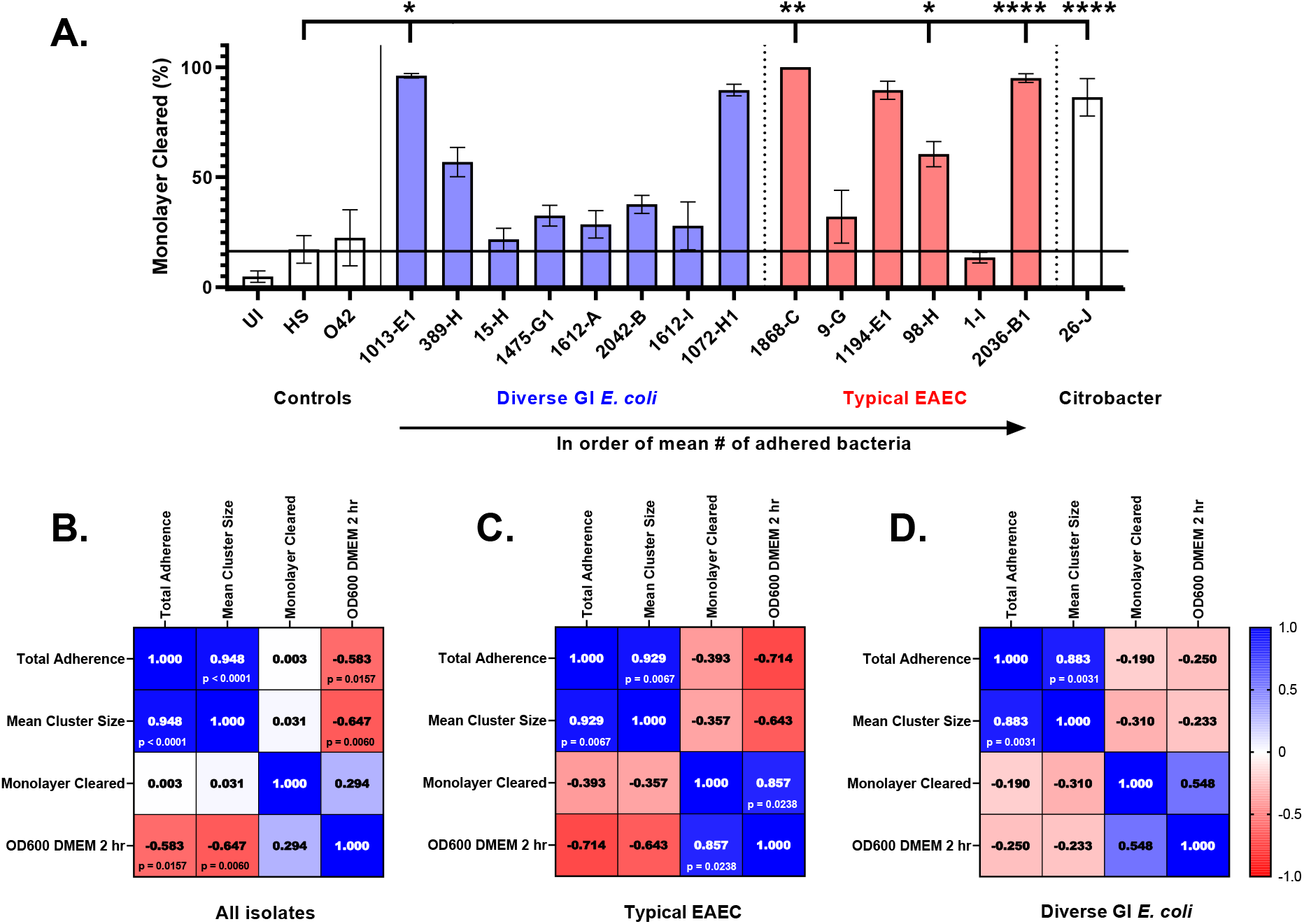
Cytotoxicity of E. coli isolates to colonoid HIEs and correlation matrices comparing bacterial adherence, bacterial turbidity as a surrogate for inoculum size, and monolayer clearance. For cytotoxicity assays, TC202 monolayers were infected with *E. coli* for 6 hours at an MOI = 25, fixed, stained, and imaged. The area of the monolayer remaining post-infection (A) was measured using FIJI/ImageJ. Experiments where >50% of the uninfected control wells had lost >10% of the monolayer were excluded. Five strains (typical EAEC strains 1868-C, 98-H, and 2036-B1; diverse GI *E. coli* isolate 1013-E1; and *Citrobacter* strain 26-J) had significantly less monolayer remaining post-infection compared to HIEs infected with control strain HS. Mean is plotted with error bars depicting SEM. Kruskal Wallis test for non-parametric data with Dunn’s multiple comparison test. All other comparisons were not significant. An increased MOI of 25 (in comparison to adherence experiments with an MOI of 10) was chosen for this experiment to increases our ability to detect bacteria capable of clearing the monolayer. For some strains which cleared the monolayers almost in totality, for example, 1868-C, the decreased MOI = 10 allowed adherence to be accessed without a majority of the monolayer destroyed. To determine if cytotoxicity was related to bacteria’s ability to grow, adhere, or aggregate, we compared several measures the clearance data. We compared the turbidity (OD600) of each isolate subcultured for 2 hours (mimicking the procedure prior to infection), as well as total adherence and mean bacterial cluster size. Data from all *E. coli* isolates and the controls HS and 042 were compared using Spearman’s correlation test for non-parametric data (B). Additionally, typical EAEC strains (all isolates and control strain 042) were analyzed together (B) and all diverse GI isolates were analyzed together (D). Significant comparisons are labeled on the figure and are as follows: (B) All isolates: total adhered bacteria and mean cluster size (r(19) = [0.948], p < [0.0001]; total adhered bacteria and bacterial growth (r(15) = [- 0.583], p = [0.0157]); mean cluster size and bacterial growth (r(15) = [-0.647], p = [0.0060]); (C) Typical EAEC: total adhered bacteria and mean cluster size (r(5) = [0.929], p = [0.0067]); monolayer clearance and bacterial growth (r(5) = [0.857], p = [0.0238]); and (D) Diverse GI *E coli*: Total adhered bacteria and mean cluster size r(7) = [0.883], p = [0.0031]). All other comparisons were not significant.

To better visualize unique features, bacterial clusters were then sorted into size categories (e.g. 3-10, 11-50, etc.) and the mean number of bacteria in each size category graphed (Figure 3E). Amongst the typical EAEC, we observed that as the total number of adherent bacteria increased, clusters >1000 became the overwhelming majority and smaller clusters become less common. Among the diverse GI *E. coli*, isolates 1013-E1, 389-H, 15-H, and 1475-G1 displayed remarkably low adherence and did not form clusters of >50 bacteria; these strains are noticeably poor at adherence compared not only to their fellow isolates, but also to non-pathogenic control HS. However, some of the strains (most notably 1072-H1) displayed adherence and aggregation in similar ranges as known diarrheagenic *E. coli* including EPEC, EIEC, and ETEC.

### Cytotoxic Potential

The cytotoxic potential of *E. coli* strains was determined by measuring the extent of epithelial peeling from the intestinal monolayer that was present at 6 hours post infection at an MOI of 25 bacteria:HIE cell (Figure 4A). Five strains (3 typical EAEC strains, 1 diverse GI *E. coli* isolate, and the *Citrobacter* isolate) caused significant HIE detachment from the chambered slides when compared to HIEs infected with the control strain HS. This was a particularly striking observation considering that the prototype EAEC strain 042 did not cause significant cell clearance in this colonoid model. Three of the six (50%) typical EAEC strains (1868-C, 98-H, and 2036-B1; p= 0.0026, p = 0.0193, and p = <0.0001 respectively), and 1/8 (12.5%) of the diverse *E. coli* isolates (1013-E1; p = 0 .0203); and the sole *Citrobacter* strain (26-J; p = <0.0001) caused significant HIE damage when compared to the non-pathogenic control HS. Isolate 1072-H1 also showed a highly cytotoxic trend. Interestingly, monolayer disruption and detachment did not correlate with total number of adhered bacteria, mean cluster size, nor growth in cell culture media when assessed for all isolates (Figure 4B). When we looked only at EAEC strains (Figure 4C), we found a positive correlation between monolayer clearance and growth in cell culture media (DMEM/F12) (r(5) = [0.857], p = [0.0238]) which was not true for the diverse GI isolates (Figure 4D).

### Virulome Characterization of EAEC and other diverse GI *E. coli*

To better appreciate the factors that may be driving the observed adherence and cytotoxicity phenotypes, we seqenced all *E. coli* isolates using a combination of whole genome PacBio sequencing and performed a “virulome” analysis^37^ to examine the presence of known or putative virulence factors present in each strain. This was achieved by aligning and comparing the WGS data to a panel of *Enterobacteriaceae* genes coding for pathogenic properties, including those related to adherence, toxin production, iron uptake, and capsule biosynthesis (Figures 5, 6, 8, and Sup. Figure 1). The phylogroup, sequence type, serotype, and fimH type were also determined from genomic data (Sup. Table 1), ^37^ and a phylogenetic tree was generated to show how relatedness of the isolates to each other and to already characterized *E. coli* strains (Figure 9). When the virulomes of our *E. coli* set were examined for pathogenic factors, they displayed an assortment of such genes, including adherence fimbriae, autotransporters, outer membrane proteins, and capsule elements (Figures 5, 6, 8, and Sup. Figure 1). None of the strains in our study carried the characteristic virulence factors^38^ for EPEC (eaeA, bfpA), ETEC (LT, ST), EHEC (Stx1, Stx2), or EIEC (ipA III, ipaIV).

**Figure 5.**
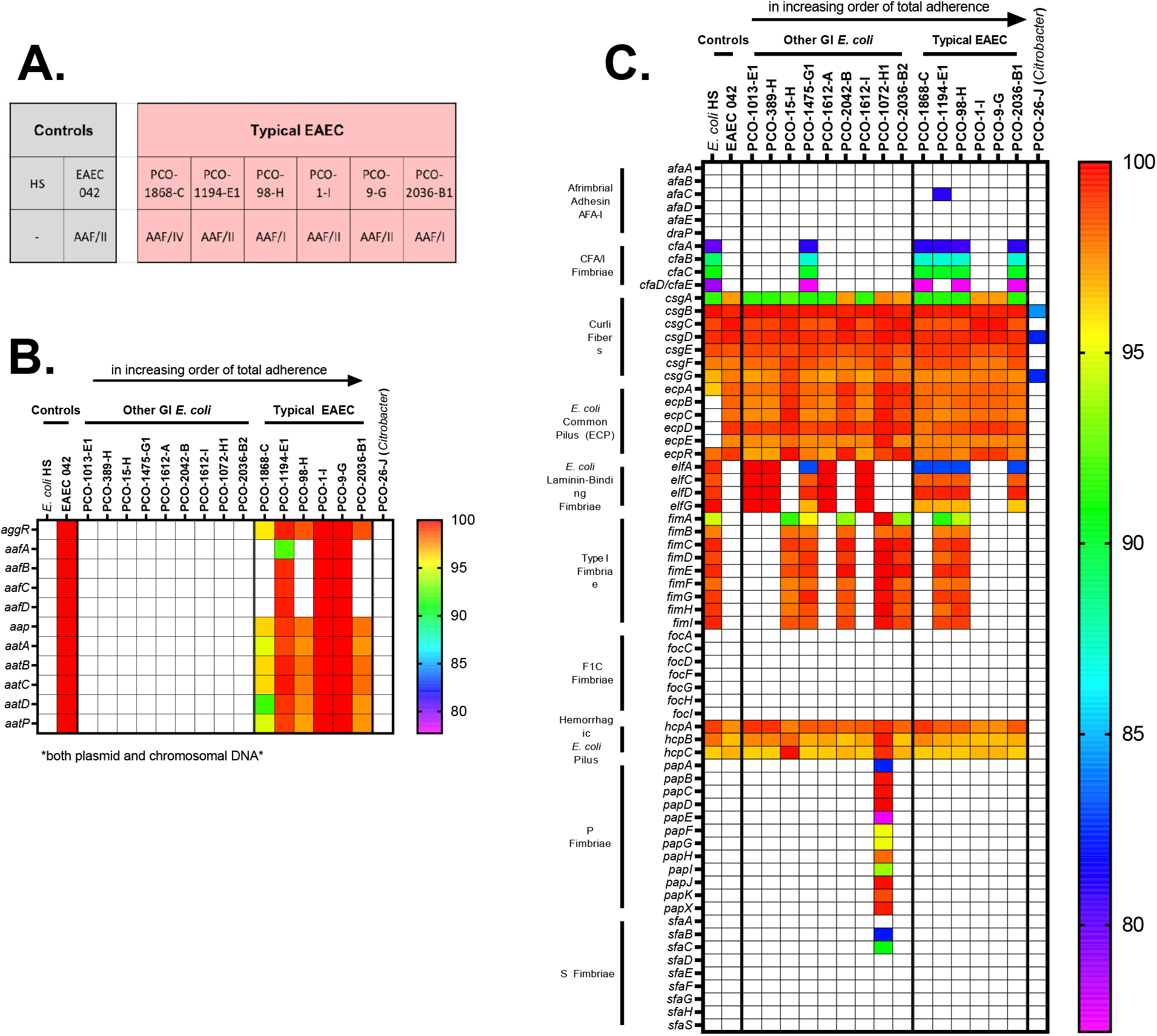
Adherence-related virulome heat maps for strains in this study. We determined the Aggregative Adherence Fimbriae (AAF) type of each of our typical EAEC strains (A) using whole genomic data. Genes associated with EAEC (including the individual genes which comprise the 5 AAF cassette variants) are depicted in (B). Additional adherence genes of *E. coli* are characterized in (C). Color represents the percentage of shared base pairs with the reference sequence matched to the key to the right of each heatmap.

We began by searching each isolate for the genes necessary for the aggregative phenotype and other genes with known association with EAEC (Figure 5B) using the genome of EAEC 042 as the reference standard. Using this data, the six clinical isolates were found to carry *aggR*, dispersin (*aatA*) and its transporters (*aatB, aatC, aatD*, and *aatP*), as well as a complete aggregative adherence cassette. The AFF adhesin types^20^ found are shown in Figure 5A. Isolates 1-I and 9-G shared 100% similarity with the AAF/II cassette of EAEC 042 while strain 1194-E1 had some divergence. Two strains, 98-H and 2036-B1, contained AAF/I cassettes while 1868-C carried the AAF/IV variant. We investigated all strains for the CS22-like gene cluster recently identified^20^ and found that no strains contained this gene cluster. Both typical EAEC and the diverse GI isolates carried several adhesins and fimbriae (Figure 5C). All isolates were found to contain the *E. coli* common pilus (ECP) and curli fibers while some isolates also carried CFA/I Fimbriae and Type I Fimbriae. Interestingly, 1072-H1 is the only isolate harboring P fimbriae, an adherence factor for UroPathogenic *E. coli* (UPEC)^39^.

Unexpectedly, we noticed a relationship between metal acquisition genes and bacterial adherence. As depicted in Figure 6A, when the isolates were aligned in order from least to greatest by total adherence, it became apparent that the 10 strains with the greatest adherence (all 6 EAEC strains and the next 4 diverse GI *E. coli* strains) carry multiple metal utilization genes. All 10 of these strains carried the iron/manganese transporter genes (*sitB, sitC,* and *sitD)* and salmochelin siderophore genes (*iroB, iroC, iroD, iroE, iroE,* and *iroN*) while none of the isolates with the poorest adherence carried these genes. In addition to these 9 shared genes, several of the isolates also carried additional genes needed to utilize metals. Using a Spearman test, we found that the number of metal genes per strain held a significant positive correlation with total bacterial adherence (r(15) = [0.5580], p = [0.02]) (Figure 6B). This relationship suggests a role for metal genes in aiding or facilitating bacterial adherence.

**Figure 6.**
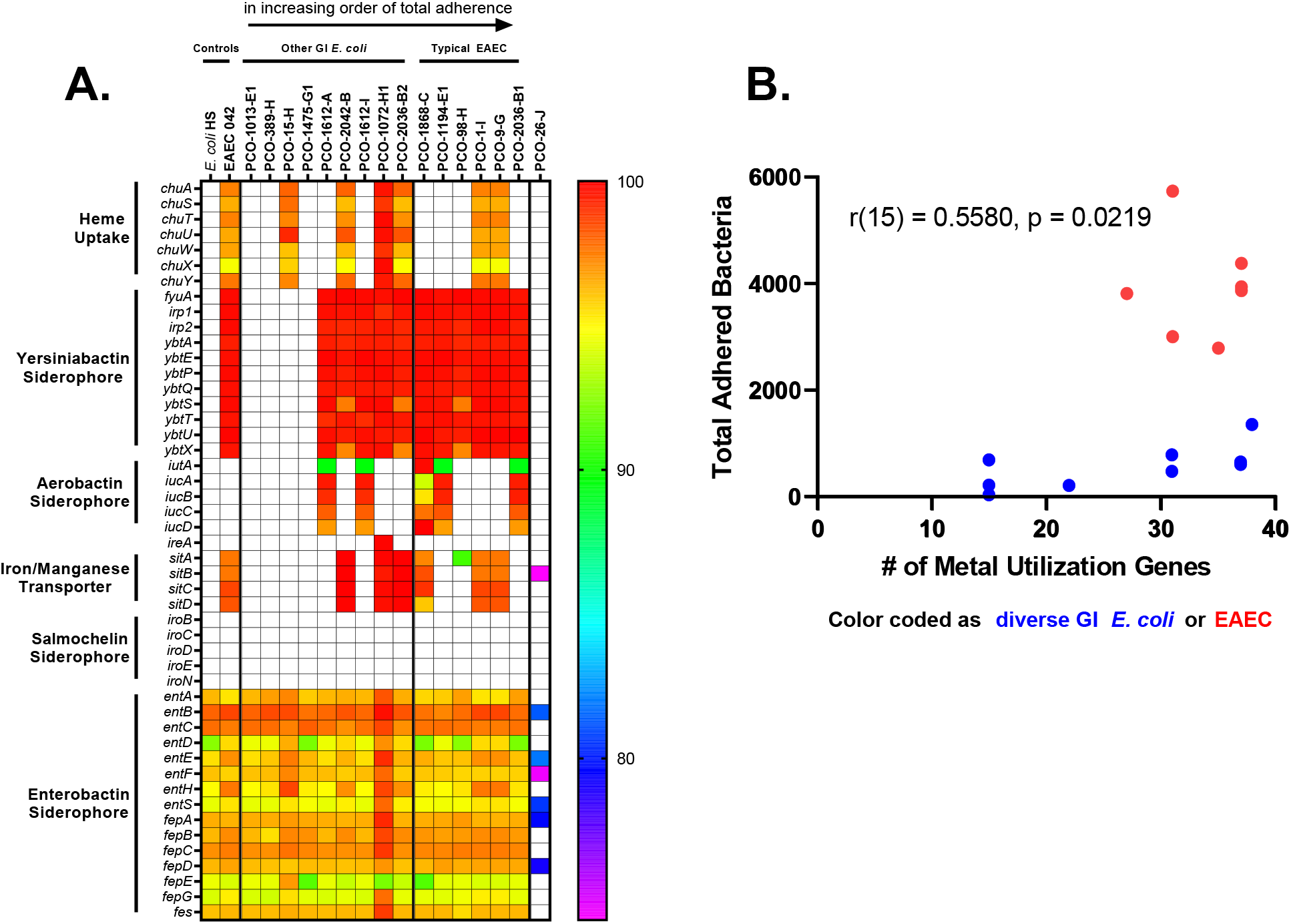
Metal acquisition and utilization virulome. Heat map visualization (A) of *E. coli* virulence factors for usage of iron and other metals revealed a positive correlation between total adherence and the number of metal genes carried (B). This analysis included all *E. coli* strains isolated from patient samples and excluded the *Citrobacter freudii* strain 26-J.

Hypothesizing that the formation of biofilm-like aggregates could decrease susceptibility to antibiotics, we tested antibiotic susceptibility (Figure 7, Sup. Figures 1 and 2) and characterized their carriage of antibiotic resistance genes. Both EAEC isolates and diverse GI *E. coli* contained a plethora of genes conferring resistance to clinical antibiotics (Sup. Figure 1). 100% of strains in both categories were sensitive to aminoglycosides, carbapenems, and nitrofurantoin while some isolates showed resistance to β-lactams, cephalosporins, monobactams, fluoroquinolones, and sulfonamides. The presence of genes linked to antibiotic resistance did not always correspond to tested resistance to that antibiotic class.

**Figure 7.**
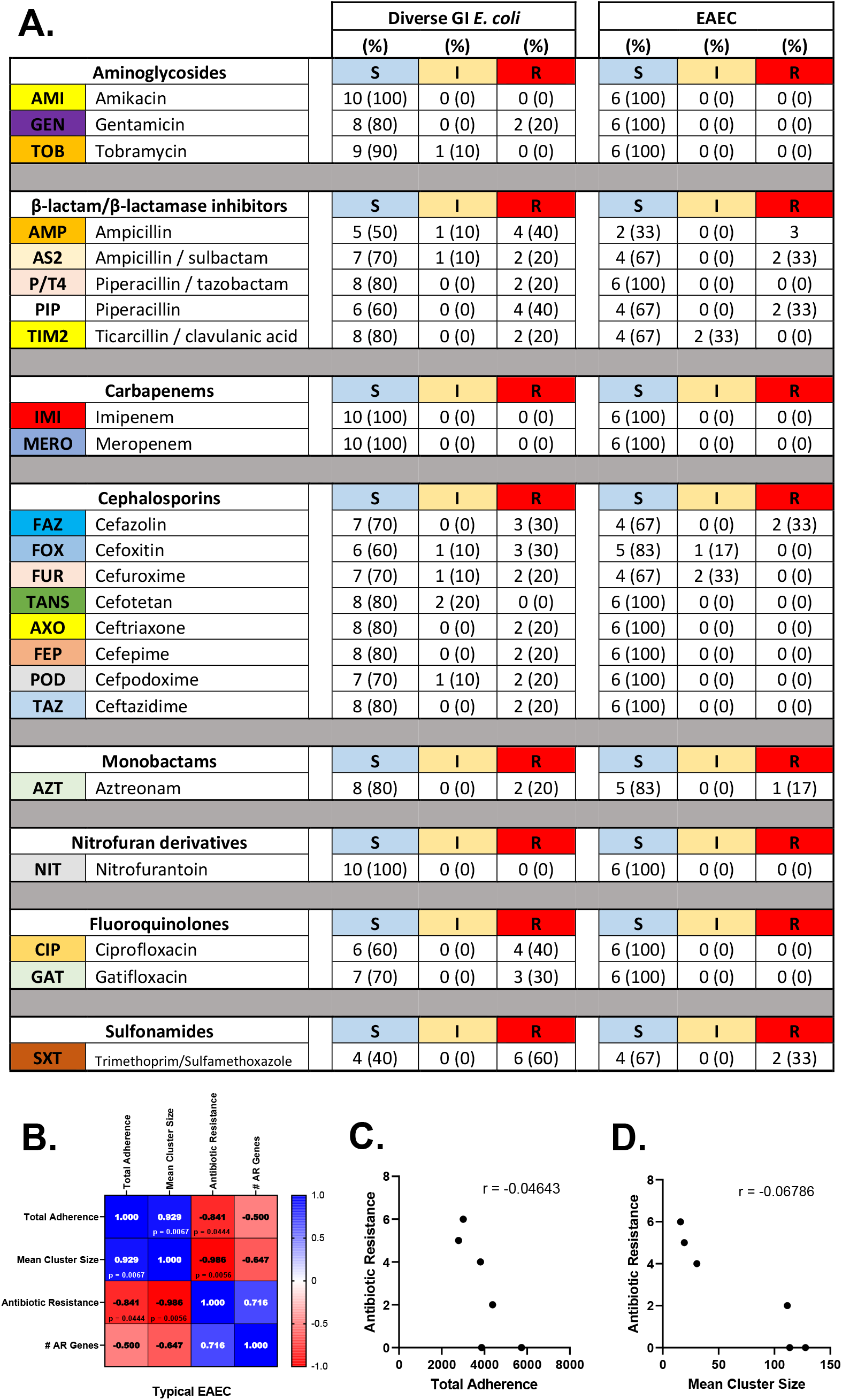
Antimicrobial susceptibility of EAEC and diverse GI E. coli. Susceptibility to a panel of clinical antibiotics was tested using and cut-offs determined using the breakpoints as recommended by CLSI M100, 32nd Edition, 2022 (A). S = sensitive to antibiotics listed, I = intermediate sensitivity, R = resistant to the antibiotics listed. Spearman’s correlation test was used to compare data from multiple assays for EAEC (B-D) and the diverse GI *E. coli* (data not shown). Within the diverse GI *E. coli*, no correlations were significant. For EAEC, the number of antibiotics each strain was resistant to (above “antibiotic resistance”) negatively correlated with both total adherence (C) and mean cluster size (D). p-values for all significant correlations are listed (B). An unlisted p-value indicates a non-significant comparison.

However, carriage of at least 3 mutations conferring resistance to fluoroquinolones (Supplemental Figure 2) did correspond to MIC testing. For each strain, we compared the number of clinical antibiotics they were resistant to (Figure 7B “Antibiotic Resistance”), the number of antibiotic resistance genes they carried, total adherence, and mean cluster size. For diverse GI *E. coli* as a group, there were no significant correlations between these features (data not shown). However, for EAEC, we unexpectedly found that greater antibiotic resistance was negatively correlated with both total adherence and mean cluster size (Figure 7B-D).

In regards to carriage of known virulence factors, all the isolates belonged to either Group 2, 3, or 4 capsule gene groups (Figure 8A) with no pattern discerning between EAEC and other diverse GI isolates. All isolates, both EAEC and other strains, carried at least one autotransporter (Figure 8B) including several which are implicated in adherence or auto-aggregation (e.g. *ang43*, *cah*, *aida,* and *FimH).* With respect to putative or known toxins (Figure 8C), the diverse GI *E. coli* only contained hemolysin E (which incidentally we also identified in strain HS). Five of the 6 typical EAEC isolates carried *pic* and both subunits of enterotoxin-1 (EAST-1) (s*et1A* and *set1B).* In addition to these virulence factors, strain 1194- E1 also carried *sat* and 2036-B1 also carried *pet*. Interestingly isolate 98-H lacked all of these accessory virulence factors associated with pathogenic EAEC. All the diverse GI *E. coli* lacked known EAEC associated genes (including aggregative adherence genes, *pic, pet, set, and sat*).

**Figure 8.**
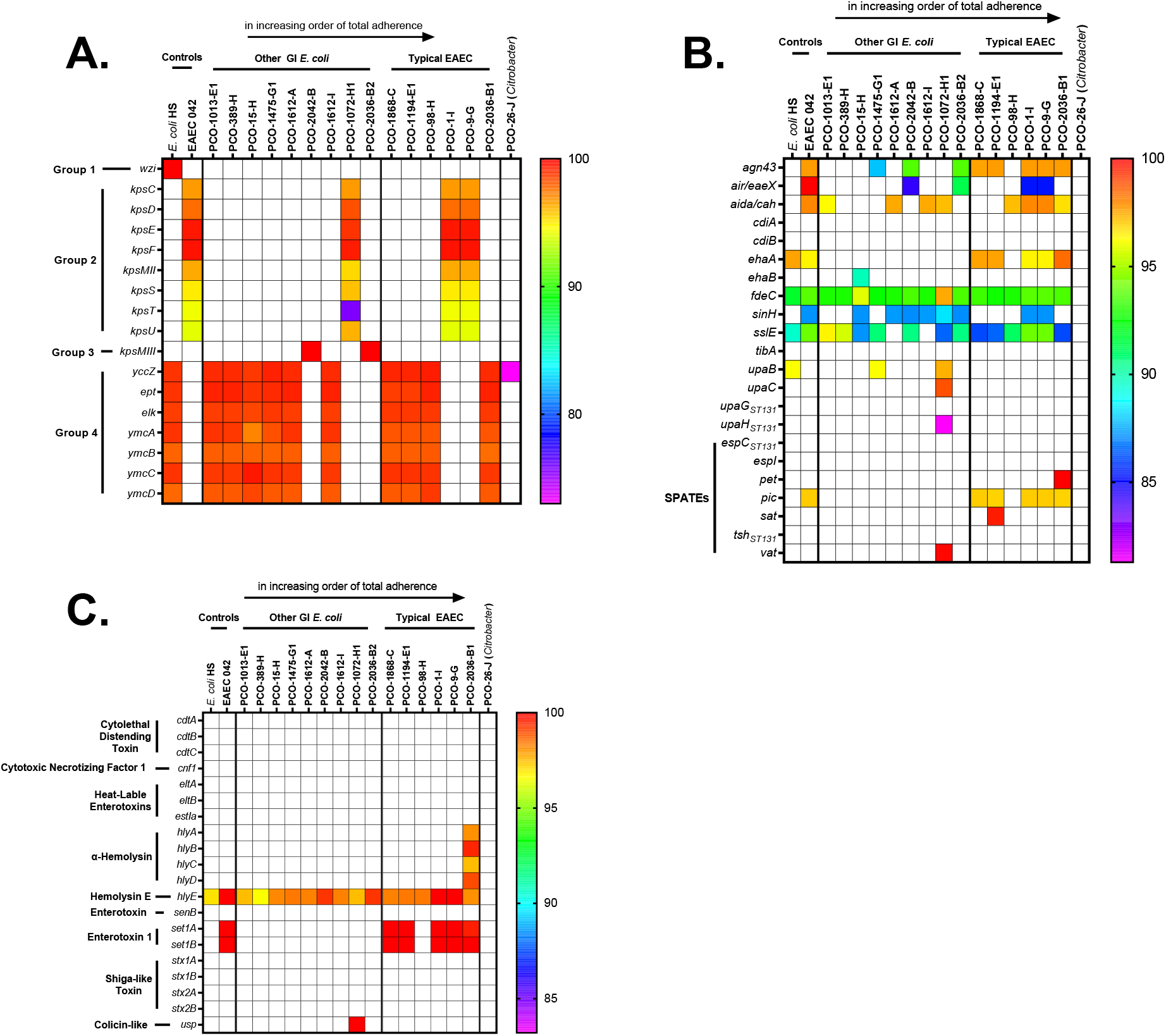
Virulome heat maps for other known virulence factors of E. coli. Capsule elements (A), Autotransporters (B) including Serine Protease Autotransporters of *Enterobacteriaceae* (SPATES), and secreted toxins (C).

Perhaps the most striking of all features was the noticeable lack of any known toxins in isolate 1013-E1 despite both being similar in cytotoxicity to the most toxic EAEC *E. coli* in our entire data set. In contrast, strain 1072-H1 contains a gene that encodes USP (colicin-like), a protein with demonstrated genotoxic effects^40^, which is often found in UPEC strains. Strain 1013-E1 is also the *E. coli* with the poorest adherence, even relative to non-pathogenic HS, of all the *E. coli* accessed in this study, a microcosm of the general decoupling of cytotoxicity from adherence capability. The lack of evidence for the presence of known cytotoxic genes that could be considered to destroy a colonoid monolayer in this strain suggests there are novel ways in which *E. coli* can damage the intestinal epithelium that remain to be discovered. Further studies will focus on identifying this possible mechanism.

The similarly of each of our isolates to well-characterized strains, using a phylogenetic analysis, shed some light on several of our diverse GI *E. coli* isolates (Figure 9). All typical EAEC strains matched to phylogroups with known association with EAEC (the B1 phylogroup is associated with both EAEC and commensal strains and the D phylogroup is associated with both EAEC and extraintestinal pathogenic *E. coli* (ExPEC)^37^). We found that the diverse GI *E. coli* in our analysis are overall genetically related to both pathogenic (B2, D, B1) and traditionally non-pathogenic (A, G, B1) phylogroups.

**Figure 9.**
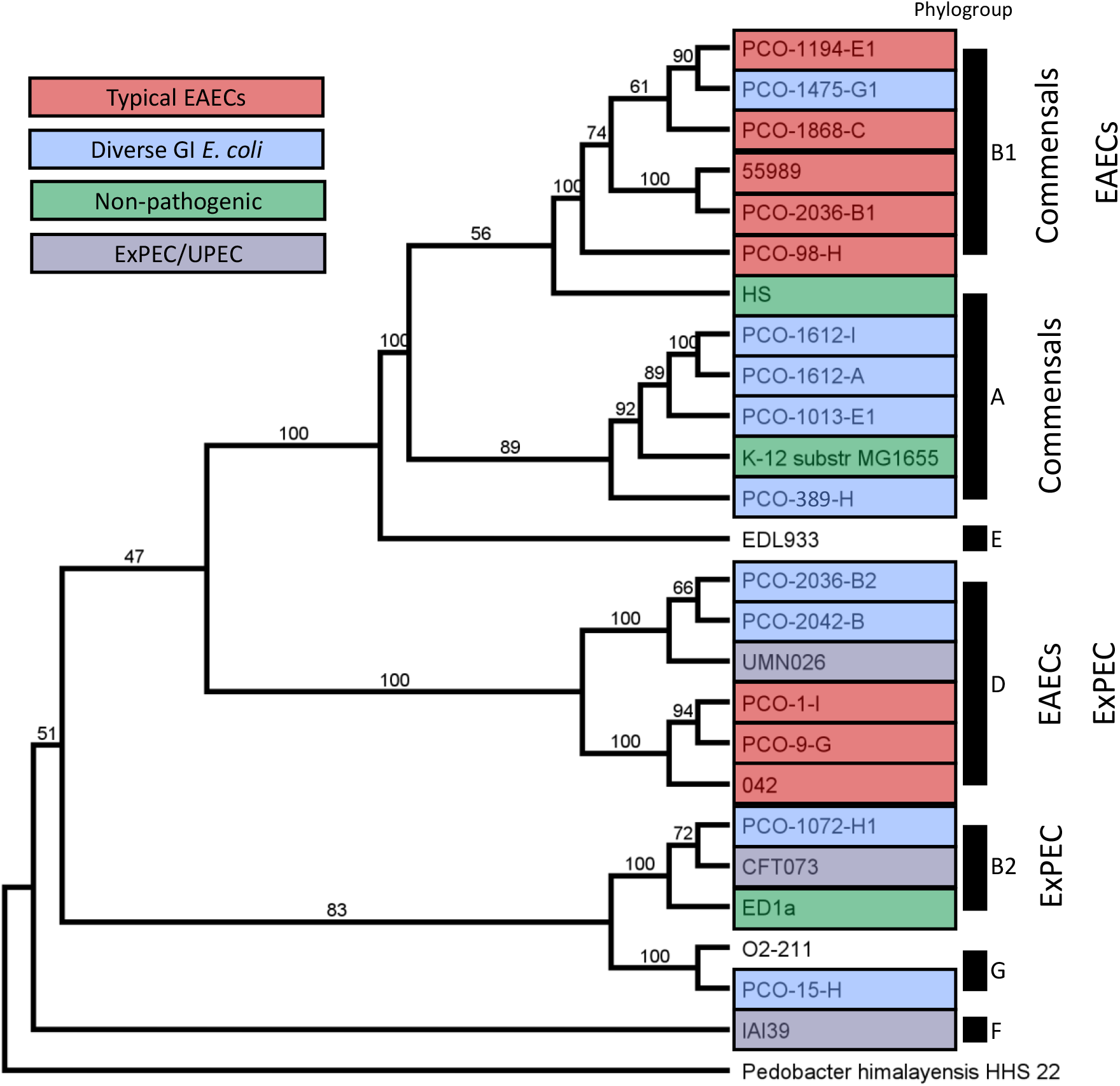
Phylogrouping of strains in this study. Phylogenetic tree created using Multi-Locus Sequence Analysis (MLSA) with autoMLST software. Similarly of each of our isolates to well-characterized strains shed some light on several of our diverse GI *E. coli* isolates. Branches are labeled with percent consensus from 1,000 UltraFast BootStrap replicates. EDL933 is an EHEC isolate. O2-211 is avian pathogenic *E. coli* (APEC). ED1a was isolated from a healthy patient.

Several of the diverse GI *E. coli* strains shared factors currently associated with UPEC and other ExPEC. Two of the diverse GI *E. coli* isolates, 1072-H1 and 2042-B, are closely related to ExPEC strains. Strain 1072-H1 clustered with the B2 phylogroup UPEC strain CFT073 and also carried P fimbriae (Figure 5) which is important for UPEC colonization^41^. Two EAEC isolates, as well as the protype EAEC strain 042, were found to be in phylogroup D along with isolate 2036-B2 and isolate 2042-B which clustered with UMN026^42^, an ExPEC strain.

Despite phylogroup A being predominately comprised of commensal strains, curiously, isolates 1612-A and 1612–I of the A phylogroup shared characteristics with strain VR50, a UPEC strain isolated from an asymptomatic bacteriuria. Isolates 1612-A and 1612-I uniquely carried sinH, yersiniabactin, and aerobactin (Figure 6B) – a rare combination in phylogroup A only observed in VR50 which carries yersiniabactin and aerobactin^37^.

Three of the non-EAEC *E. coli* strains are members of EAEC-associated phylogroups (strains 1475-G1 of phylogroup B1 and 2036-B2 and 2042-B of phylogroup D). Four of the diverse GI strains are in phylogroup A which is mostly associated with commensal *E. coli* suggesting that these strains are either non-pathogenic, opportunistic pathobionts, or were previously commensal *E. coli* which later acquired virulence capability.

## DISCUSSION

The role of EAEC as a causal agent of diarrhea in patients with cancer is unclear. This is in part due to the sensitivity and specificity of methods used for detection, travel history, rates of colonization, genetic heterogeneity of strains, degree of immunosuppression, antibiotic therapy and existing co-morbidities that can cause diarrhea such as mucositis, neutropenic enterocolitis, use of laxatives, hyperosmotic enteral feeding or medication induced diarrhea including chemotherapy in this patient population.

In this study, we extend our previous observations on diarrheagenic *E. coli* in patients with cancer in whom EAEC is the third most common bacterial enteropathogen identified after *C. difficile* and EPEC when using sensitive NAATs^43^. Using novel, stem-cell derived HIEs as a model of infection, and whole genome sequencing, we contrasted the bacterial adherence patterns, burden of infection, and genetic virulence factor profile of clinical EAEC isolates, reference strains belonging to the major diarrheagenic *E. coli* pathotypes, and a diverse group of intestinal *E. coli*. We found that NAATs overestimate the frequency of EAEC in cancer patients with diarrhea, that EAEC clinical isolates are strong biofilm formers in HIEs and demonstrate significant heterogeneity in HIE cytotoxicity and that the presence of metal utilization genes in this collection of EAEC and diverse *E. coli* is associated with enhanced bacterial adherence to colonoid HIEs. We also show ETEC LT, a pathogen that is associated with small bowel colonization and secretory diarrhea avidly binds to colonoid HIEs. In addition to these findings, there is a noticeable lack of positive correlation between either total adherence or aggregation and colonoid cytotoxicity whereby strains with strong adherence are not necessarily more cytotoxic than strains with less adherence.

Consistent with previous studies, EAEC was isolated as a pure culture in only 6/29 (20%) of stool samples with EAEC identified by NAAT. A variety of reasons could explain this including antibiotic use prior to sample collection resulting in NAAT positive results but non-viable organisms in culture and/or colonization with EAEC with a burden that is below the threshold for bacterial culture detection methods. In this study, the likelihood of isolating and successfully characterizing *E. coli* strains was significantly higher in patients in who were severely immunocompromised following HSCT and is likely a reflection of carrying higher bacterial loads as has been shown for EPEC in this population^32^. This is particularly important when testing was done in patients with diarrhea that is due to other causes such as mucositis or chemotherapy in whom EAEC was incidentally detected by NAAT. Given the large shifts in microbiota in cancer patients receiving antibiotics, and dominance of *Enterobacteriaceae*, it is also plausible that EAEC could represent false positive results due to probe cross reactivity to genes with similarity to *aggR*. Additionally, disagreement between NAAT results and recovery of EAEC in culture can occur due to loss of the PAA plasmid carrying *aggR* due to passaging (which seems plausible for strain 1475-G1 because on whole genomic sequencing, the backbone resembled reference EAEC strains). Nevertheless, our findings raise some concern on the sensitivity and specificity of NAATs in this patient population.

All reference pathotypes including ETEC LT adhered to colonoid HIEs. Robust aggregative phenotype was observed for the reference EAEC strain 042 and for all 6 clinical EAEC isolates. Of interest, 6 of the 9 diverse *E. coli* that did not carry virulence factors classically associated with diarrheagenic *E. coli* also adhered to the human intestinal colonoids in equal number as prototype strains of EIEC, ETEC, EHEC, and EPEC and disturbed the HIE monolayer. A few strains formed small aggregative clusters that in the absence of molecular testing for *aggR* could have been categorized as EAEC if phenotype was the sole criteria used for EAEC definition. One such isolate, 1072, also caused significant destruction of the intestinal monolayer and was found to be a UPEC strain carrying the P fimbriae confirming the importance of molecular testing to differentiate typical EAEC from atypical EAEC and the known overlap in phenotypes between UPEC and EAEC. Of note, this strain is in the B2 phylogroup which have a high carriage rate of virulence factors^37^.

A subset of the diverse GI *E. coli* in this series are likely commensals (devoid of virulence factors that showed little or no adherence to HIE. However, strain 1013 also caused significant damage to the HIEs despite no clear explanation within the virulome results and absolute poor performance (i.e., the worst colonoid binder) of all the *E. coli* in this study. While there is no clear definition of commensalism in *E. coli* for humans, it is generally assumed that strains from healthy patients with no association with disease (such as the prototype strain HS^33^) act as harmless inhabitants of the GI tract. However, in patients at risk for infection, it is possible that these *E. coli* strains can become pathobionts and cause human disease. It is also possible that, although more physiologic than other model systems, colonoids are lacking factors that may be present as disease drivers or accessory adherence factors *in vivo*.

EAEC frequently co-exists with other enteropathogens ^27^ and can sometimes cause bacteremia^44, 45^ whether EAEC participates in consortium with other pathogens in the GI tract is unknown. The identification of clinical of ExPEC and cytotoxic *E. coli* that do not fit classical pathotypes in patients with diarrhea is concerning as this could lead to bacteremia from translocation in patients with dysbiosis, intestinal inflammation or loss of barrier function following exposure to antibiotics and chemotherapy^7, 46,47,48^.

EAEC pathotype classification remains challenging because the relationship between the virulence factors harbored and the diseases caused remains unclear. For example, EAST-1, which we identified in 5 of our 6 EAEC isolates, has been implicated in fluid secretion^49, 50^, but its sufficiency to cause diarrhea is still controversial^51^. We found that EAEC isolate 2036-B1 showed unique cytotoxic potential, clearing >95% of the monolayer, and was the sole strain capable of damaging the monolayer even when the MOI was decreased fivefold (data not shown). We note that this isolate is the only one in our study to carry alpha hemolysin A shown in the toxin heatmap (Figure 8C). Also, noteworthy, isolate 98-H differed from other EAEC strains in that it lacks both EAST-1 and pic, yet still damaged the HIE as determined by >50% clearance. A recent finding indicates that diarrheal disease is associated with EAEC of sequence type ST40 while asymptomatic infection is higher among ST31 strains^8^. Interestingly, none of our EAEC isolates from this patient population belonged to these sequence types (Sup. Table 1).

Additionally, it remains unclear how the formation of aggregative adherence *in vitro* relates to symptomatic disease in human patients. Though all five variants of AAF have been demonstrated to form aggregative adherence to Hep-2 cells and AAF/I and AAF/II have additionally been tested in other transformed intestinal cell lines^8^, only adherence to HIEs had only been demonstrated with AAF/II^15, 31^. Here we confirm that EAEC strains with AAF/I, AAF/II, and AAF/IV all form robust aggregative adherence on human intestinal enteroids. Furthermore, adherence number and pattern are indistinguishable between these three AAF types.

The presence of metal acquisition genes in EAEC isolates has been previously reported^20, 52^ and iron has been shown to be important in the formation of EAEC biofilms^53^. Our findings suggest a role for metals in EAEC adherence and aggregation on the human intestinal epithelium. Though many of these genes are primarily thought of as iron-utilization genes, siderophores can interact with an array of metals. The yersiniabactin siderophore carried by all of our EAEC isolates, as well as the best adhering non-EAEC diverse GI *E. coli*, can be used to acquire zinc^54^. The presence of zinc at physiological levels has been shown to decrease EAEC’s ability to form biofilms and adhere to intestinal epithelial cells^55^. Zinc supplementation has been shown to be of benefit in pediatric diarrhea, but trials using zinc as a complementary treatment for patients undergoing chemotherapy have seen mixed results^56^. Targeting metal utilization of EAEC through small molecule inhibitors, supplementation, or vaccination could provide new treatment options.

Resistance to one or more antibiotics has been identified in up to 91.8% of clinical EAEC strains ^10, 49, 57^. In our study, we identified genes conferring resistance to numerous antibiotic drug classes in both EAEC and the diverse GI *E. coli* (Sup. Figure 1) including extended spectrum beta lactamases CTX-M 15 and OXA-1. Although most infections due to EAEC are self-limited and do not require treatment, severe infections may require antibiotic therapy. The use of antibiotics can further select for resistant organisms, limit treatment options and lead to dysbiosis and *Enterobacteriaceae* microbiome dominance^6, 7^ which in the right setting can be a precursor to bacteremia. Whether colonization with biofilm forming EAEC or carriage of adherent *E. coli* serve as a reservoir for MDR genes that can be exchanged with ExPEC or other Gram-negative rods in cancer patients is unknown. Alternative therapies are needed to prevent and treat *E. coli* infections in this patient population. The presence of shared virulence factors in EAEC and ExPEC strains suggests that a common target could be used for the development of a vaccine against both pathotypes.

The breadth of *E coli* pathogenesis variance is becoming increasingly complex as evidenced by the relatively small number of strains studied here. Whole genome analysis of clinical *E. coli* strains is a powerful tool and is yielding promising insights into pathogenicity, treatment targets and importantly vaccine candidates^37^. The creation of accessible biobanks with curated and annotated epidemiologic data, clinical information from healthy controls and patients coupled with powerful sequencing technologies, and new models of adhesion and cytotoxicity, such as the HIEs described here allow the opportunity to re-define *E. coli* pathotypes with greater diagnostic accuracy and into more clinically relevant groupings. As recent publications have demonstrated^8, 20^, refining or restructuring heterogeneous pathotypes may require us to abandon dearly held paradigms. Future studies are likely to redefine and design new categorization schemes to better inform clinical treatment and to facilitate development of new therapeutic agents and vaccines. We plan to characterize additional clinical isolates by whole genome sequencing and using HIEs to identify additional virulence factors associated with adhesion and cytotoxicity, to improve the resolution of diagnostic tests, identify new treatment targets for small molecular inhibitors of bacterial adherence, or serve as vaccine candidates.

## METHODS

### Human Intestinal Enteroid Culture

The human intestinal enteroid line TC202 was established from health patient crypts (transverse colon of patient 202) and maintained in CMFG+ media as previously described^58^. Colonoids benefited from the addition of an increased concentration of Wnt during maintenance and passaging. HIEs were plated as monolayers on Matrigel (Corning #356231) coated plates (Greiner Bio One #543979 and Corning 96 well plates) and differentiated 4-5 days prior to infection.

### Isolation of *E. coli* Strains

Fecal samples were collected from patients with cancer and diarrhea in whom infectious cause was suspected, stools were placed on Cary Blair transport media and studied for the presence of enteropathogens using the BioFire FilmArray® gastrointestinal panel platform in the clinical microbiology laboratory as standard of care. This study was reviewed by the University of Texas MD Anderson Cancer Center Institutional Review Board and deemed to be exempt from IRB review. Residual stool samples were transported to the research laboratory and streaked onto a MacConkey agar plate. Ten coliform colonies were selected from each plate and stored as agar stabs as described previously^59^. Each stored agar stab from this collection was streaked onto MacConkey agar and from each, 10 colonies were chosen and streaked onto EMB agar. Individual green and black colonies (indicative of *E. coli*) were selected and streaked again on EMB agar to isolate individual strains. Single colonies were grown overnight in LB broth and stored as both a frozen stock and a peptone stab. MALDI TOF mass spectrometry was used to identify the species of each isolate and species was later confirmed by genomic analysis.

### Control and reference strains

Isolate HS^60^, a non-pathogenic *E. coli* strain which does not cause diarrheal disease in human adults and does not carry known genes associated with the diarrheagenic pathotypes^33^ was used as a negative control. EAEC 042 (kindly gifted by Dr. James Nataro), first isolated from a child in Peru^61^, has been studied extensively and was used as a positive control for typical EAEC. Additionally, prototype strains of each diarrheagenic *E. coli* pathotype were used for comparison: EIEC NCDC 909-51^62^ (ATCC 23520), EHEC 86-24^63^ (kindly gifted by Dr. Alfredo Torres), EPEC E2348/69^33^, and ETEC LT H10407^64^ (ATTC 35401).

### Bacterial Adherence Assays

Bacteria were grown overnight shaking in LB media at 37°C and then subcultured for 3 hours^65^ in DMEM. HIE monolayers were infected at an MOI of 10 by dilution of the subculture in differentiation media and incubated 6 hours in static conditions at 37C, 5% CO2 as previously described^15, 65^. Media was removed and slides were fixed and stained with a Hema3 kit and imaged using the Nikon 100X Plan / 1.25 NA objective. Three images were taken per well and multiple wells were imaged for each condition.

Bacterial adherence was counted and quantified as previously described^32^. A bacterial cluster was defined as a group of bacteria which were visibly touching one another or with a gap of <0.5 micron between them (to allow for close interactions via biofilm extracellular polysaccharides and/or capsule proteins).

### Cytotoxicity Screen

HIE monolayers differentiated 4-5 days were infected with *E. coli* at an MOI=10:1 for 6 hours. Media was removed and slides were fixed 5 minutes with methanol, stained with a Hema3 kit, and allowed to dry. Images of the slides were taken, and the area of the monolayer in each well measured using ImageJ. The percentage of the monolayer cleared post-infection was calculated using the following formula 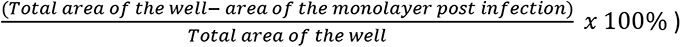 as previously described^32^

### Library Preparation, Sequencing, and Genetic Analyses

Frozen bacterial pellets were thawed and the DNA was extracted using Phenol Chloroform. Bacterial genomic DNA samples was mechanically sheared to ∼15kb using Covaris g-TUBE. Fifteen of the samples in the study was prepped using SMRTbell Template preparation kit 1.0 (Pacific Biosciences, Menlo Park, CA, USA), while a single sample (PCO-2036-B2) was prepped using the newer SMRTbell Express Template preparation kit 2.0. The libraries further prepared according to the PacBio standard protocol. The purified ligation productions were then pooled and size-selected on Sage Science BluePippin. The 15 samples were run as a pool of 4-plex on PacBio Sequel I with 20 hrs movie time and the single sample was run as part of another set of 20-plex samples on PacBio Sequel II 8M SMRT cell with a 30 hrs movie time.

Assembly of the raw PacBio subreads was performed using Canu (version 1.8^66–68)^ with an estimated genome size of 4.8Mb and a minimum read length 2Kb. Assembly contigs were subsequently polished by aligning the raw subreads to them using the PacABio tool blasr^69^. The resulting alignments were then analyzed using PacBio’s tool "arrow" to yield corrected contigs.

Following annotation, assembled contigs were used as queries in megaBLAST (blast 2.9.0)^70^ to search against both a database of *E. coli* virulence factors curated by our lab^37^ and the Comprehensive Antimicrobial Resistance Database (CARD) protein homology database (version 3.1.4)^71^, with a culling and HSPS limit of 2 (“-culling_limit 2” and “-max_hsps 2”, respectively) but otherwise using default settings in Geneious Prime 2022.0.2. To focus on functional hits, truncated genes and HSPSs that were less than 90% the length of the subject were omitted. Percent identity was extracted and used to produce heatmaps using GraphPad Prism version 9.3.1 for Windows, GraphPad Software, San Diego, California USA, www.graphpad.com.

Phylotypes of assembled genomes were determined bioinformatically using in-house^37^ and open-source software^72, 73^. Sequence type was determined using pubMLST^74^ and the Center for Genomic Epidemiology’s (CGE) MLST 2.0 software (version 2.0.4)^75, 76^. Finally, serotype and FimH type was determined using CGE’s SerotypeFinder 2.0 (version 2.0.1; database version 1.0.0)^77^ and FimTyper 1.0 (version 1.0)^78^, respectively. All settings were default unless otherwise noted.

Phylogenetic tree was generated from Multi-Locus Sequence Analysis (MLSA) performed by the Automated Multi-Locus Species Tree (autoMLST)^79^ software with default settings set to produce a concatenated alignment with 1,000 replicates with IQ-TREE Ultrafast Bootstrap analysis. Resulting tree was visualized in Geneious Prime 2022.0.2. For comparison, strains HS (accession: CP000802), ED1a (accession: CU928162), K12 substrain MG1655 (accession: CP014225), 042 (accession: FN554766), 55989 (accession: CU928145), EDL933 (accession: CP008957), IAI39 (accession: CU928164), UMN026 (accession: CU928163), CFT073 (accession: AE014075), and O2-211 (accession: CP006834) were used. All PacBio Contigs from clinical isolates were submitted to NCBI (BioProject PRJNA812607). BioSample accession IDs are listed in supplementary table 1.

### Statistical analyses

For statistical comparisons, either one-way ANOVA (with Dunnett’s multiple comparisons test) or (Kruskal Wallis test with Dunn’s multiple comparison test) was selected based on the normality of the data set. In all figures, **** denotes p = 0.0001 - 0.001, *** denotes p = 0.001 - 0.01, ** denotes p = 0.01 to 0.05, *denotes p ≥ 0.05.

## ACKNOWLEDGEMENTS

We gratefully acknowledge all participants of the TMC-GCID team. A list of TMC-GCID members can be found on the center’s website: https://gcid.research.bcm.edu/. Thank you to Dr. Alfredo Torres and Dr. James Nataro for sharing *E. coli* strains with our laboratory. Imaging for this project was supported by the Integrated Microscopy Core at Baylor College of Medicine with funding from NIH. This work was funded by grants from the National Institutes of Health (NIH): U19AI144297 (Integrated Genomics of Mucosal Infections) and U19AI116497 (Human Gastrointestinal Biomimetics for Enteric Bacterial Infections).

## SUPPLEMENTAL

**Supplemental Table 1.**
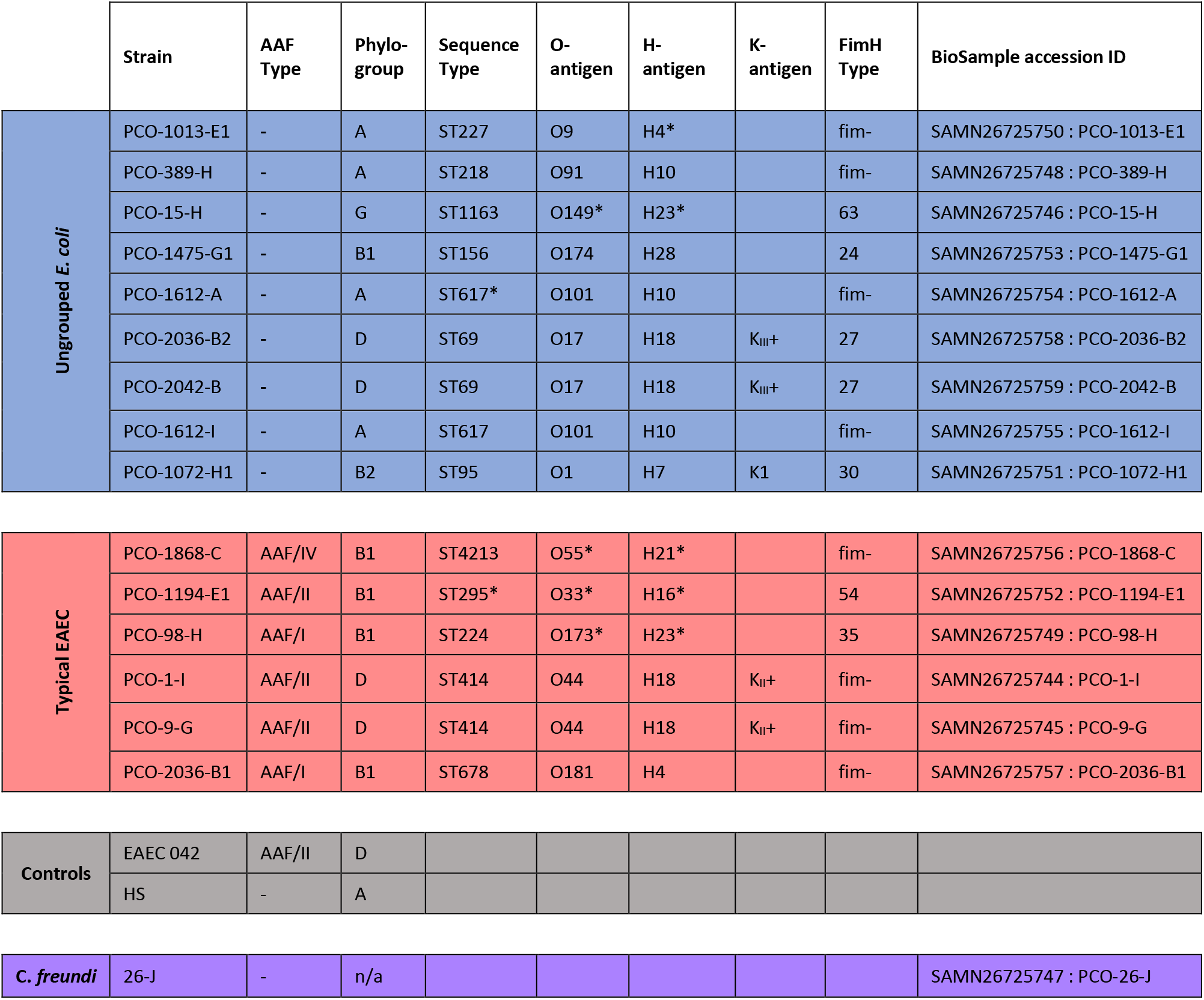
E. coli typing and phenotype summaries for strains in this study. Aggregative adherence to multiple substrates were characterized and are summarized here. Former *E. coli* typing methods (phylogroup, sequence type, O-antigen, H-antigen, K-antigen, and FimH type) and AAF type were determined from sequence data. * indicates imperfect match.

**Supplemental Figure 1.**
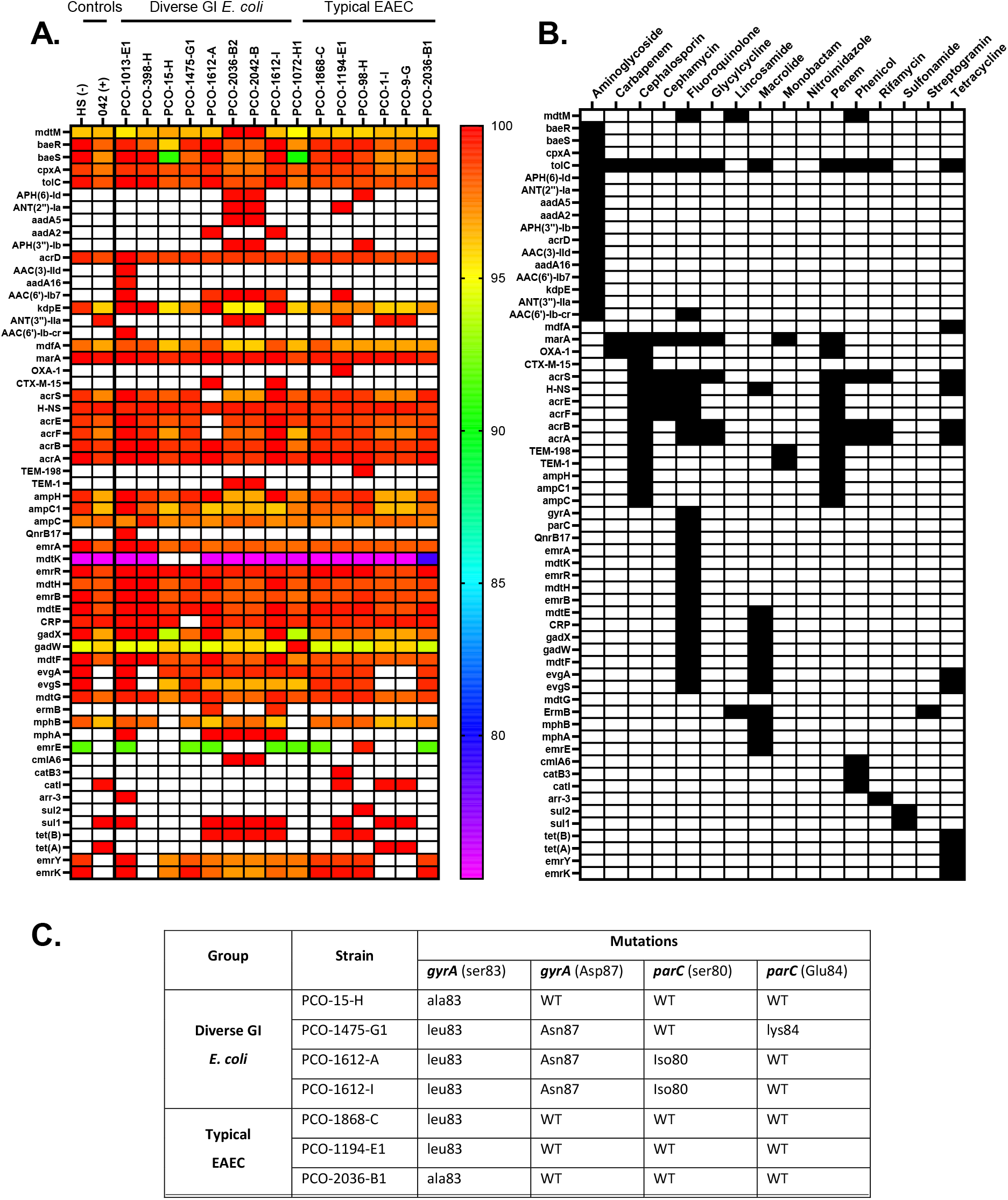
Clinical antibiotic resistome. (A) Heatmap depicting the similarity of each gene for resistance to clinical antibiotics mapped to the reference sequence. The antibiotic class each gene corresponds to is matched in (B) which is based on the Comprehensive Antimicrobial Resistance Database (CARD)’s Antibiotic Resistance Ontology (ARO) classifications. Point mutations in *gyrA* and *parC* can confer resistance to fluoroquinolones but are not apparent in heat map presentation. *GyrA* and *parC* sequences from each strain were aligned to the reference sequence to look for specific mutations conferring fluoroquinolone resistance (C). Three of the EAEC strains had mutations at ser83 of *gyrA.* We also observed the well characterized mutations at ser83 and asp87 of *gyrA* and ser80 and glu84 of *parC* which confer resistance to fluoroquinolones^80–82^ among the diverse GI *E. coli*. All other isolates had wildtype amino acids at these gene locals. No mutations were observed at Gln106 of *gyrA* in these strains. Additionally, 1194-E1 also had a frameshift mutation in *parC* beginning at the 104^th^ amino acid.

**Supplementary Figure 2.**
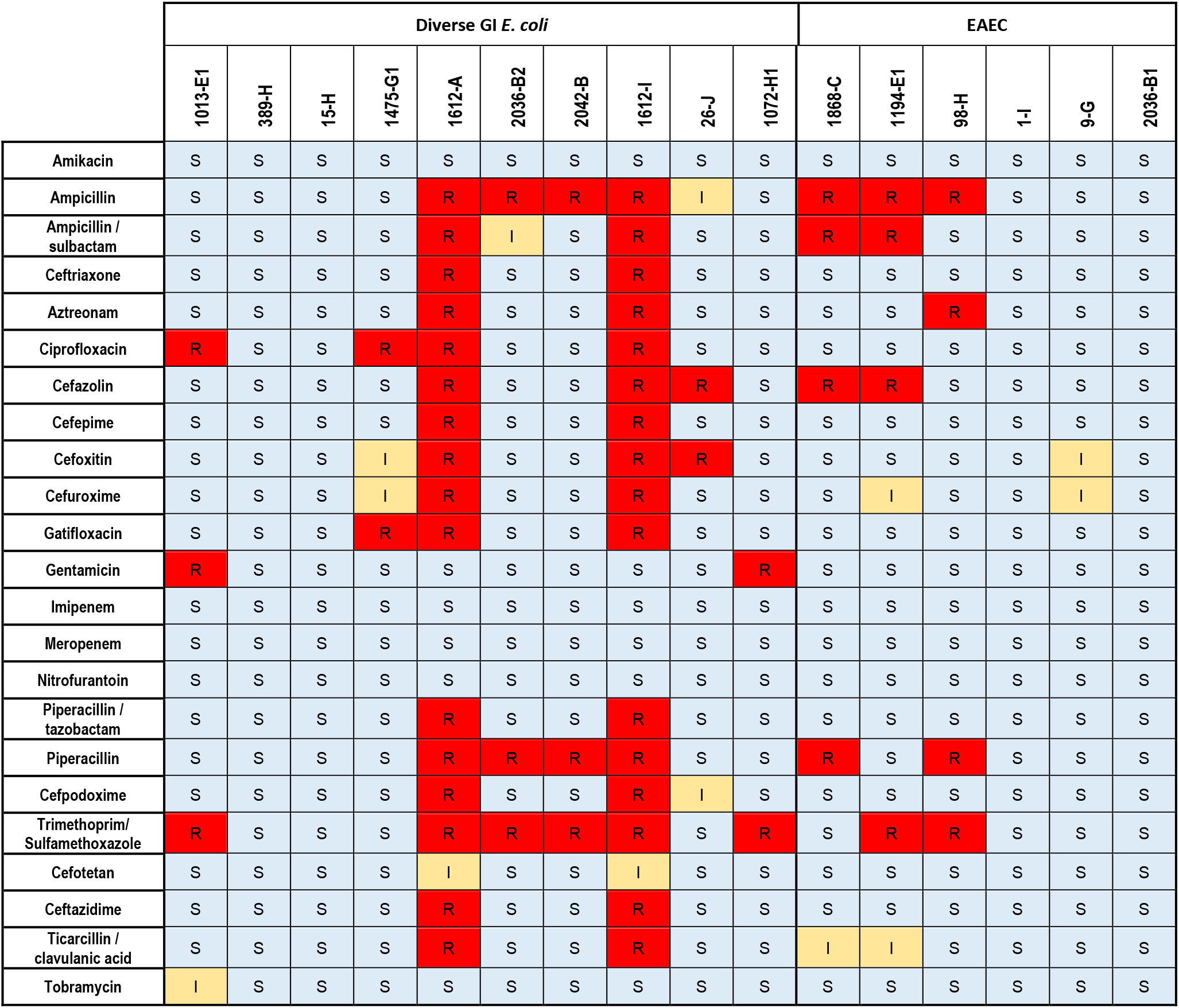
Antimicrobial susceptibility results by each isolate. Sensitivity or resistance was determined based on the tested minimum inhibitory concentration (MIC) compared to the breakpoints recommended by CLSI M100, 32nd Edition, 2022. S = sensitive to antibiotics listed, I = intermediate sensitivity, R = resistant to the antibiotics listed.

## REFERENCES

1. Zembower, T. R. Epidemiology of infections in cancer patients. Cancer Treatment and Research vol. 161 (2014).

2. Nosari, A. et al. Infections in haematologic neoplasms: autopsy findings. Haematologica 76, 135– 140 (1991).

3. Homsi, J. et al. Infectious complications of advanced cancer. Support. Care Cancer 8, 487–492 (2000).

4. Chang, H. Y. et al. Causes of death in adults with acute leukemia. Med. 55, 259–68 (1976).

5. Ubeda, C. et al. Vancomycin-resistant Enterococcus domination of intestinal microbiota is enabled by antibiotic treatment in mice and precedes bloodstream invasion in humans. J. Clin. Invest. 120, 4332–4341 (2010).

6. Hueso, T. et al. Impact and consequences of intensive chemotherapy on intestinal barrier and microbiota in acute myeloid leukemia : the role of mucosal strengthening. Gut Microbes 12, 1–17 (2020).

7. Conn, J. R., Catchpoole, E. M., Runnegar, N., Mapp, S. J. & Markey, K. A. Low rates of antibiotic resistance and infectious mortality in a cohort of high-risk hematology patients: A single center, retrospective analysis of blood stream infection. PLoS One 12, 1–13 (2017).

8. Ellis, S. J. et al. Identification and characterisation of enteroaggregative Escherichia coli subtypes associated with human disease. Sci. Rep. 10, 1–12 (2020).

9. Jensen, B. H., Olsen, K. E. P., Struve, C., Krogfelt, K. A. & Petersen, A. M. Epidemiology and clinical manifestations of enteroaggregative escherichia coli. Clin. Microbiol. Rev. 27, 614–630 (2014).

10. Okeke, I. N. et al. Heterogeneous virulence of enteroaggregative Escherichia coli strains isolated from children in Southwest Nigeria. J. Infect. Dis. 181, 252–260 (2000).

11. Nataro, J. P. et al. Heterogeneity of enteroaggregative escherichia coli virulence demonstrated. J. Infect. Dis. 171, 465–468 (1995).

12. 12. Nataro, J. P., et al. Heterogeneity of Enteroaggregative " Escherichia coli " Virulence Demonstrated in Volunteers Published by : Oxford University Press Stable URL : http://www.jstor.org/stable/30132045 JSTOR is a not-for-profit service that helps scholars, researchers, and. 171, 465–468 (2016).

13. Asea, A., Kaur, P. & Chakraborti, A. Enteroaggregative Escherichia coli: An emerging enteric food borne pathogen. Interdiscip. Perspect. Infect. Dis. 2010, (2010).

14. Steiner, T. S., Lima, A. A. M., Nataro, J. P. & Guerrant, R. L. Enteroaggregative Escherichia coli produce intestinal inflammation and growth impairment and cause interleukin-8 release from intestinal epithelial cells. J. Infect. Dis. 177, 88–96 (1998).

15. Rajan, A. et al. Novel segment- and host-specific patterns of enteroaggregative escherichia coli adherence to human intestinal enteroids. MBio (2018) doi:10.1128/mBio.02419-17.

16. Jiang, Z. D. et al. Genetic susceptibility to enteroaggregative Escherichia coli diarrhea: Polymorphism in the interleukin-8 promotor region. J. Infect. Dis. 188, 506–511 (2003).

17. Mohamed, J. A. et al. A novel single-nucleotide polymorphism in the lactoferrin gene is associated with susceptibility to diarrhea in North American travelers to Mexico. Clin. Infect. Dis. 44, 945–952 (2007).

18. Mohamed, J. A. et al. Single nucleotide polymorphisms in the promoter of the gene encoding the lipopolysaccharide receptor CD14 are associated with bacterial diarrhea in US and Canadian travelers to Mexico. Clin. Infect. Dis. 52, 1332–1341 (2011).

19. Nataro, J. P. et al. Patterns of adherence of diarrheagenic Escherichia coli to HEp-2 cells. Pediatr. Infect. Dis. 6, 829–831 (1987).

20. Boisen, N. et al. Redefining enteroaggregative escherichia coli (Eaec): Genomic characterization of epidemiological eaec strains. PLoS Negl. Trop. Dis. 14, 1–19 (2020).

21. Pereira, A. L., Silva, T. N., Gomes, A. C. & Arajo, A. C. Diarrhea-associated biofilm formed by enteroaggregative Escherichia coli and aggregative Citrobacter freundii: A consortium mediated by putative F pili. BMC Microbiol. 10, (2010).

22. Lang, C. et al. Novel type of pilus associated with a Shiga-toxigenic E. coli hybrid pathovar conveys aggregative adherence and bacterial virulence. Emerg. Microbes Infect. 7, (2018).

23. Dias, R. C. B. et al. Analysis of the Virulence Profile and Phenotypic Features of Typical and Atypical Enteroaggregative Escherichia coli (EAEC) Isolated From Diarrheal Patients in Brazil. Front. Cell. Infect. Microbiol. 10, (2020).

24. Dudley, E. G. et al. An IncI1 plasmid contributes to the adherence of the atypical enteroaggregative Escherichia coli strain C1096 to cultured cells and abiotic surfaces. Infect. Immun. 74, 2102–2114 (2006).

25. Čobeljić, M. et al. Enteroaggregative Escherichia coli associated with an outbreak of diarrhoea in a neonatal nursery ward. Epidemiol. Infect. 117, 11–16 (1996).

26. Harrington, S. M., Dudley, E. G. & Nataro, J. P. Pathogenesis of enteroaggregative Escherichia coli infection. FEMS Microbiol. Lett. 254, 12–18 (2006).

27. Chattaway, M. A., et al. Investigating the link between the presence of enteroaggregative Escherichia coli and infectious intestinal disease in the United kingdom, 1993 to 1996 and 2008 to 2009. Eurosurveillance 18, (2013).

28. Garcia, P. G., Silva, V. L. & Diniz, C. G. Occurrence and Antimicrobial Drug Susceptibility Patterns of Commensal and Diarrheagenic Escherichia Coli in Fecal Microbiota from Children with and without Acute Diarrhea. J. Microbiol. 49, 46–52 (2011).

29. Nüesch-Inderbinen, M. T., Hofer, E., Hächler, H., Beutin, L. & Stephan, R. Characteristics of enteroaggregative Escherichia coli isolated from healthy carriers and from patients with diarrhoea. J. Med. Microbiol. 62, 1828–1834 (2013).

30. Kaper, J. B., Nataro, J. P. & Mobley, H. L. T. Pathogenic Escherichia coli. Nat. Rev. Microbiol. 2, 123–140 (2004).

31. Rajan, A. et al. Enteroaggregative E . coli Adherence to Human Heparan Sulfate Proteoglycans Drives Segment and Host Specific Responses to Infection. Plos Pathog. 16, 1–27 (2020).

32. Olvera, A. et al. Enteropathogenic Escherichia coli Infection in Cancer and Immunosuppressed Patients . Clin. Infect. Dis. 72, (2020).

33. Levine, M. M. et al. Escherichia Coli Strains That Cause Diarrhœa But Do Not Produce Heat-Labile or Heat-Stable Enterotoxins and Are Non-Invasive. Lancet 311, 1119–1122 (1978).

34. In, J. et al. Enterohemorrhagic Escherichia coli Reduces Mucus and Intermicrovillar Bridges in Human Stem Cell-Derived Colonoids. Cmgh 2, 48–62.e3 (2016).

35. Regua-Mangia, A. H. et al. Genotypic and phenotypic characterization of enterotoxigenic Escherichia coli (ETEC) strains isolated in Rio de Janeiro city, Brazil. FEMS Immunol. Med. Microbiol. 40, 155–162 (2004).

36. Okhuysen, P. C. & Dupont, H. L. Enteroaggregative Escherichia coli (EAEC): A Cause of Acute and Persistent Diarrhea of Worldwide Importance. 202, 503–505 (2010).

37. Clark, Justin. Maresso, A. Comparative Pathogenomics of Escherichia coli : Polyvalent Vaccine Target Identification Through Virulome Downloaded from http://iai.asm.org/ on May 11, 2021 by guest. (2021) doi:10.1128/IAI.00115-21.

38. Aranda, K. R. S., Fagundes-Neto, U. & Scaletsky, I. C. A. Evaluation of Multiplex PCRs for Diagnosis of Infection with Diarrheagenic Escherichia coli and Shigella spp . J. Clin. Microbiol. 42, 5849–5853 (2004).

39. Lane, M. C. & Mobley, H. L. T. Role of P-fimbrial-mediated adherence in pyelonephritis and persistence of uropathogenic Escherichia coli (UPEC) in the mammalian kidney. Kidney Int. 72, 19–25 (2007).

40. Armstrong, G. D. Uropathogenic escherichia coli colicin-like usp and associated proteins: Their evolution and role in pathogenesis. J. Infect. Dis. 208, 1539–1541 (2013).

41. Melican, K. et al. Uropathogenic Escherichia coli P and Type 1 Fimbriae Act in Synergy in a Living Host to Facilitate Renal Colonization Leading to Nephron Obstruction. Plos Pathog. 7, 2–13 (2011).

42. Kwak, M. J., Kim, M. S., Kwon, S. K., Cho, S. H. & Kim, J. F. Genome sequence of Escherichia coli NCCP15653, a group D strain isolated from a diarrhea patient. Gut Pathog. 8, 1–7 (2016).

43. Chao, A. W. et al. Clinical features and molecular epidemiology of diarrheagenic Escherichia coli pathotypes identified by fecal gastrointestinal multiplex nucleic acid amplification in patients with cancer and diarrhea. Diagn. Microbiol. Infect. Dis. 89, 235–240 (2017).

44. Boll, E. J. et al. Emergence of enteroaggregative Escherichia coli within the ST131 lineage as a cause of extraintestinal infections. bioRxiv 11, 1–13 (2018).

45. Mandomando, I. et al. Escherichia coli st131 clones harbouring AGGR and AAF/V fimbriae causing bacteremia in Mozambican children: Emergence of new variant of FIMH27 subclone. PLoS Negl. Trop. Dis. 14, 1–21 (2020).

46. Crawford, J., Dale, D. C. & Lyman, G. H. Chemotherapy-Induced Neutropenia: Risks, Consequences, and New Directions for Its Management. Cancer 100, 228–237 (2004).

47. Russo, F. et al. The effects of fluorouracil, epirubicin, and cyclophosphamide (FEC60) on the intestinal barrier function and gut peptides in breast cancer patients : an observational study. BMC Cancer 13, 1–11 (2013).

48. Green, S. I. et al. Murine model of chemotherapy-induced extraintestinal pathogenic Escherichia coli translocation. Infect. Immun. 83, 3243–3256 (2015).

49. Zamboni, A., Fabbricotti, S. H., Fagundes-Neto, U. & Scaletsky, I. C. A. Enteroaggregative Escherichia coli Virulence Factors Are Found to Be Associated with Infantile Diarrhea in Brazil. J. Clin. Microbiol. 42, 1058–1063 (2004).

50. Savarino, S. J., Fasano, A., Robertson, D. C. & Levine, M. M. Enteroaggregative Escherichia coli elaborate a heat-stable enterotoxin demonstrable in an in vitro rabbit intestinal model. J. Clin. Invest. 87, 1450–1455 (1991).

51. Ruan, X., Crupper, S. S., Schultz, B. D., Robertson, D. C. & Zhang, W. Escherichia coli expressing EAST1 toxin did not cause an increase of cAMP or cGMP levels in cells, and no diarrhea in 5-day old gnotobiotic pigs. PLoS One 7, (2012).

52. Okeke, I. N., Scaletsky, I. C. A., Soars, E. H., Macfarlane, L. R. & Torres, A. G. Molecular Epidemiology of the Iron Utilization Genes of Enteroaggregative Escherichia coli. J. Clin. Microbiol. 42, 36–44 (2004).

53. Alves, J. R. et al. Iron-limited condition modulates biofilm formation and interaction with human epithelial cells of enteroaggregative Escherichia coli (EAEC). J. Appl. Microbiol. 108, 246–255 (2010).

54. Behnsen, J. et al. Siderophore-mediated zinc acquisition enhances enterobacterial colonization of the inflamed gut. Nat. Commun. 12, (2021).

55. Medeiros, P. et al. The micronutrient zinc inhibits EAEC strain 042 adherence, biofilm formation, virulence gene expression, and epithelial cytokine responses benefiting the infected host. Virulence 4, 624–633 (2013).

56. Hoppe, C. et al. Zinc as a complementary treatment for cancer patients: a systematic review. Clin. Exp. Med. 21, 297–313 (2021).

57. Estrada-Garcia, T. & Navarro-Garcia, F. Enteroaggregative Escherichia coli pathotype: A genetically heterogeneous emerging foodborne enteropathogen. FEMS Immunol. Med. Microbiol. 66, 281–298 (2012).

58. Winnie Y. Zou, Sarah E. Blutt, Sue E. Crawford, Khalil Ettayebi, Xi-Lei Zeng, Kapil Saxena, Sasirekha Ramani, Umesh C. Karandikar, Nicholas C. Zachos, and M. K. E. Human Intestinal Enteroids: New Models to Study Gastrointestinal Virus Infections. Physiol. Behav. 1576, 229–247 (2019).

59. Huang, D. B. et al. Virulence characteristics and the molecular epidemiology of enteroaggregative Escherichia coli isolates from travellers to developing countries. J. Med. Microbiol. 56, 1386–1392 (2007).

60. Formal, S. B., Dammin, G. J., Labrec, E. H. & Schneider, H. Experimental Shigella infections: characteristics of a fatal infection produced in guinea pigs. J. Bacteriol. 75, 604–610 (1958).

61. Nataro, J. P. . et al. Detection of an Adherence Factor of Enteropathogenic Escherichia coli with a DNA Probe. J. Infect. Dis. 152, 560–565 (1985).

62. Korst, van der. The Nederlandsch Tijdschrift voor Geneeskunde and the practice of medical science in The Netherlands (1857-1896); light under a bushel? Ned. Tijdschr. Geneeskd. 135, 1779 (1991).

63. Griffin, P. M. et al. Illnesses associated with Escherichia coli O157:H7 infections. A broad clinical spectrum. Ann. Intern. Med. 109, 705–712 (1988).

64. Skerman, F. J., Formal, S. B. & Falkow, S. Plasmid-associated enterotoxin production in a strain of Escherichia coli isolated from humans. Infect. Immun. 5, 622–624 (1972).

65. Nina M. Poole, Anubama Rajan, A. W. M. Human Intestinal Enteroids for the Study of Bacterial Adherence, Invasion and Translocation. Curr. Protoc. Microbiol. 50, (2018).

66. Koren, S. et al. Canu: scalable and accurate long-read assembly via adaptive k-mer weighting and repeat separation. Genome Res. 27, 722–736 (2017).

67. Koren, S. et al. De novo assembly of haplotype-resolved genomes with trio binning. Nat. Biotechnol. 36, 1174–1182 (2018).

68. Nurk, S. et al. HiCanu: Accurate assembly of segmental duplications, satellites, and allelic variants from high-fidelity long reads. Genome Res. 30, 1291–1305 (2020).

69. Chaisson, M. J. & Tesler, G. Mapping single molecule sequencing reads using basic local alignment with successive refinement (BLASR): Application and theory. BMC Bioinformatics 13, (2012).

70. Camacho, C., et al. BLAST+: architecture and applications. BMC Bioinformatics 10, 421 (2009).

71. Alcock, B. P., et al. CARD 2020: antibiotic resistome surveillance with the comprehensive antibiotic resistance database. Nucleic Acids Res. (2019) doi:10.1093/nar/gkz935.

72. Waters, N. R., Abram, F., Brennan, F., Holmes, A. & Pritchard, L. Easy phylotyping of Escherichia coli via the EzClermont web app and command-line tool. Access Microbiol. 2, (2020).

73. Beghain, J., Bridier-Nahmias, A., Le Nagard, H., Denamur, E. & Clermont, O. ClermonTyping: an easy-to-use and accurate in silico method for Escherichia genus strain phylotyping. *Microb*. Genomics 4, (2018).

74. Jolley, K. A., Bray, J. E. & Maiden, M. C. J. Open-access bacterial population genomics: BIGSdb software, the PubMLST.org website and their applications. Wellcome Open Res. 3, 124 (2018).

75. Larsen, M. V. et al. Multilocus sequence typing of total-genome-sequenced bacteria. J. Clin. Microbiol. 50, 1355–61 (2012).

76. Wirth, T. et al. Sex and virulence in Escherichia coli: an evolutionary perspective. Mol. Microbiol. 60, 1136–1151 (2006).

77. Joensen, K. G., Tetzschner, A. M. M., Iguchi, A., Aarestrup, F. M. & Scheutz, F. Rapid and Easy In Silico Serotyping of Escherichia coli Isolates by Use of Whole-Genome Sequencing Data. J. Clin. Microbiol. 53, 2410–2426 (2015).

78. Roer, L. et al. Development of a Web Tool for Escherichia coli Subtyping Based on fimH Alleles. J. Clin. Microbiol. 55, 2538–2543 (2017).

79. Alanjary, M., Steinke, K. & Ziemert, N. AutoMLST: an automated web server for generating multi- locus species trees highlighting natural product potential. Nucleic Acids Res. 47, W276–W282 (2019).

80. Aldred, K. J. et al. Role of the Water − Metal Ion Bridge in Mediating Interactions between Quinolones and Escherichia coli Topoisomerase IV. Biochemistry 53, 5558–5567 (2014).

81. Hallett, P. & Maxwell, A. Novel Quinolone Resistance Mutations of the Escherichia coli DNA Gyrase A Protein : Enzymatic Analysis of the Mutant Proteins. Antimicrob. Agents Chemother. 35, 335–340 (1991).

82. Lu, T., Zhao, X. & Drlica, K. Gatifloxacin Activity against Quinolone-Resistant Gyrase : Allele-Specific Enhancement of Bacteriostatic and Bactericidal Activities by the C-8-Methoxy Group. Anti 43, 2969–2974 (1999).

